# Drug-controlled CAR-T cells through the regulation of cell-cell interactions

**DOI:** 10.1101/2024.08.06.606454

**Authors:** Leo Scheller, Greta Maria Paola Giordano Attianese, Rocío Castellanos-Rueda, Raphaël B Di Roberto, Markus Barden, Melanie Triboulet, Sailan Shui, Elisabetta Cribioli, Anthony Marchand, Sandrine Georgeon, Hinrich Abken, Sai Reddy, Bruno E. Correia, Melita Irving

## Abstract

CAR T-cell therapy is constrained by on-target, off-tumor toxicities as well as cellular exhaustion due to chronic antigen exposure. CARs comprising small-molecule controlled switches can enhance both safety and therapeutic efficacy but are limited by the scarcity of non-immunogenic protein elements responsive to non-immunosuppressive, clinically approved drugs with favorable pharmacodynamics. Here, we combine rational design and library-based optimization of a protein-protein interaction (PPI) of human origin to develop venetoclax-controlled Drug-Regulated Off-switch PPI (DROP)-CARs. DROP-CARs enable dose-dependent release of the tumor-targeting scFv and consequent T-cell dissociation from the target tumor cell. Additionally, we present proof-of-concept for a dual DROP-CAR controlled by different small molecules, as well as for logic-gated synthetic receptors enabling STAT3 signaling. We demonstrate in vitro and in vivo function of DROP-CAR T cells and conclude that the approach holds promise for clinical application.

## Introduction

Chimeric antigen receptor (CAR)-T cell therapy has achieved durable and complete responses in a high proportion of patients with certain B-cell malignancies but relapse is problematic^1^. Moreover, extending such outcomes to non-hematological solid tumors which represent the majority of cancers^2^, remains a grand challenge, including from a safety perspective. Indeed, on-target, off-tumor toxicity of CARs directed against solid tumor antigens, which with few exceptions are also present on healthy tissues, is a priori a safety concern^3^. In addition, as more T-cell coengineering strategies and combinatorial treatments are implemented to address the immunosuppressive nature of the solid tumor microenvironment, the risk and incidence of toxicity to CAR therapy will likely increase^4^.

Most CARs currently used in the clinic are second generation (2G) designs and comprise an antigen-binding moiety, typically a single-chain variable fragment (scFv), fused to a hinge/linker, a transmembrane region, and the endodomains of one costimulatory receptor (usually CD28 or 4-1BB) and of CD3ζ to enable T-cell activation^5^. Limitations to the 2G-CAR include that (i) chronic exposure can render the engineered T cells exhausted^6^, (ii) toxicity can result from on-target reactivity against healthy tissues and (iii) CAR-T cell over-responsiveness in the face of high antigen-density and tumor burden can trigger adverse events in patients such as cytokine release syndrome^7^. A variety of approaches have been developed to improve the safety and efficacy of 2G-CAR T cells. For example, suicide-switches have been coengineered into T cells but once activated they terminate the therapy^8^. Alternatively, weaker affinity scFvs have been used to render the 2G-CARs irresponsive to low tumor antigen density as may be found on healthy tissue, but this may also limit efficacy due to heterogeneous expression levels on tumor cells^9–12^.

On- and off-switch CAR designs that allow remote control of T-cell activity levels by small molecule administration represent a promising strategy for balancing function and safety^5, 13^. In their pioneering work, the Lim lab developed the first on-switch split-CAR comprising FK506 binding protein (FKBP) domain and the T2089L mutant of the FKBP-rapamycin binding domain reliant upon rapamycin analog (AP21967)-mediated chain heterodimerization^14^. More recent efforts have shifted towards on- or off-switch CARs responsive to non-immunosuppressive small molecules with longer half-lives and/or that are more tolerable for long-term administration. Examples include on-switches based on viral protease inhibition in self-cleaving CARs by the antiviral agent grazoprevir^15, 16^, inducible CAR degradation by lenalidomide responsive zinc finger degrons^17^, and CARs built with antibody-fragments engineered so that tumor antigen binding/recognition is conditional upon the presence of a small molecule such as methotrexate^18, 19^.

While there has been remarkable progress in the field, switch-designs comprising human-derived domains (to minimize immunogenicity) that are responsive to clinically approved small molecules are rare, and existing ones have limitations. For example, CAR-T cell persistence/expansion correlates with patient responses but rapamycin inhibits antigen-induced proliferation of T cells^1^. In addition, the immunomodulatory effects of lenalidomide, including changes in cytokine production, T-cell activation, and NK-cell function^20^ can influence patient responses, and undesirable side-effects such as neutropenia, fatigue, and cardiac disorders^21^ impact tolerance can occur. The development of novel switch-designs controlled by alternative clinically approved small molecules is hence warranted.

Previously, we computationally developed an off-switch CAR coined the STOP-CAR and comprising receptor (R)- and signaling (S)-chains designed for reversible dissociation. The PPI (i.e., off-switch) made up of truncated Bcl-xL^22^ and LD3 (a human protein scaffold engrafted with critical binding residues from the BH3 domain of Bim to generate a high-affinity interface with Bcl-xL), was encoded in the endodomains of the chains and could be disrupted upon application of the small molecule A-1155463 ^23^. An important limitation to the STOP-CAR is that A-1155463 is not clinically approved. Moreover, because both chains are transmembrane, simple disruption of the PPI will not disengage the CAR-T cell from the target cell which may be a superior mechanism for controlling T-cell activity^24–27^.

Here, by rationally designing and screening an interface mutant library, we generated a PPI comprising truncated Bcl-2 and a variant of LD3 that is stable but can be efficiently disrupted by the clinically approved molecule venetoclax. We incorporated the optimized components into a <u>D</u>rug-<u>R</u>egulated <u>O</u>ff-switch <u>P</u>PI (DROP)-CAR design comprising a transmembrane S-chain that non-covalently engages via the PPI with a soluble R-domain. We reasoned that in this configuration disruption of the off-switch will physically disengage the DROP-CAR T cells from the target cells for more robust abrogation of effector functions. We validated controllable function of DROP-CAR T cells both in vitro and in vivo. We also demonstrated application for our optimized off-switch in the context of logic-gated cytokine receptor signaling. Taken together, we expand the toolbox of PPIs responsive to clinically approved small molecules which can help improve the safety and efficacy of cellular gene therapies.

## Results

### DROP-CAR design

Previously, we demonstrated that our human designer protein LD3 not only forms a high-affinity PPI with Bcl-xL^22^, but also with the structurally similar anti-apoptotic protein Bcl-2 (K_D_ = 0.8 nM)^28^. Moreover, LD3:Bcl-2 can be disrupted, albeit inefficiently, by the clinically approved and orally available small molecule venetoclax^28^, a Bcl-2 inhibitor used in the treatment of various hematological malignancies^29^. Here, we sought to optimize the LD3:Bcl-2 interface for use in an off-switch CAR design, on the one-hand maintaining its stability but on the other-hand allowing efficient disruption by venetoclax.

We began by building a construct encoding a heterodimeric DROP-CAR prototype with murine components targeting HER-2 and comprising Bcl-2:LD3 as a disruptable PPI. In comparison to the classic 2G-CAR which links tumor-antigen binding to T-cell activation in a single chain (**Figure 1a, left**), the DROP-CAR is a split-design comprising a transmembrane S-chain (with LD3 in the ectodomain and signaling components in the endodomain) that non-covalently associates with a coexpressed soluble R-domain (scFv fused to truncated Bcl-2) (**Figure 1a, right**) and can be disassembled in the presence of venetoclax which disrupts the PPI (Bcl-2:LD3) (**Figure 1b**). For cell-surface detection of the two DROP-CAR components, we fused a Strep-tag to the hinge of the S-chain and a V5-tag to the R-domain.

**Figure 1:**
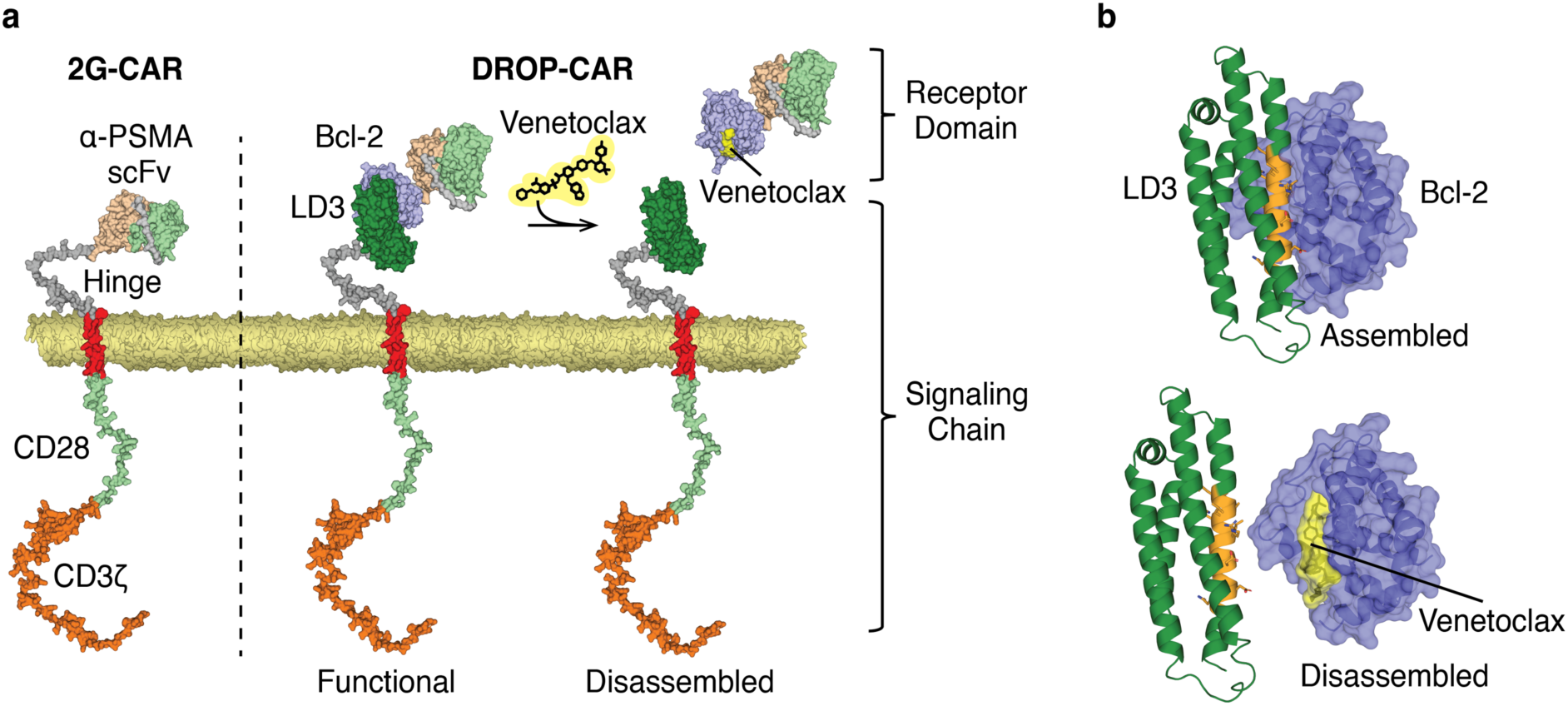
Schematic of the DROP-CAR design and venetoclax-mediated disruption of Bcl-2 and LD3. **a) Left:** Second generation (2G) CAR comprising an scFv (peach & mint green), hinge (grey), transmembrane (red), CD28 endodomain (light green) and the CD3ζ endodomain (orange). **Right**: Heterodimeric DROP-CAR comprising a Signaling (S)-chain that includes LD3 and a Receptor (R)-domain that includes the tumor targeted scFv fused to Bcl-2. In the presence of venetoclax the protein-protein interaction between Bcl-2 and LD3 is disrupted and the R-domain is released rendering the receptor disassembled and the T cells unable to respond to target antigen. **b) Top:** Model of assembled LD3:Bcl-2 complex and **Bottom:** its dissociation in response to venetoclax. The models are based on PDB 6IWB (LD3), PDB 6O0K (Bcl-2 and venetoclax), and AlphaFold2 models of receptor domains.

The DROP-CAR construct was inserted into a Cas9 expressing B3Z reporter cell line (a mouse T cell hybridoma expressing a TCR that recognizes the OVA peptide (SIIFEKL)/H-2Kb complex) in the T cell receptor beta-chain locus by homology directed repair^12^. Cell-surface expression of the assembled murine DROP-CAR was validated by dual tag-labeling and flow cytometric analysis, and we demonstrated limited disruption of the DROP-CAR after 5 h incubation with 1 µM venetoclax (**Figure S1**).

### Library design and screening to optimize LD3 for DROP-CAR use

Having demonstrated proof-of-principle for successful cell-surface assembly of the heterodimeric DROP-CAR, as well as the ability to release the R-domain upon incubation with venetoclax, we next set out to improve the PPI. We sought to better facilitate LD3:Bcl-2 disassembly by venetoclax while maintaining the stability of the S-chain:R-domain complex in its native state (i.e., the assembled DROP-CAR in the absence of venetoclax).

We began by using the Rosetta modeling suite to calculate changes in Gibbs free energy (ΔΔG) stemming from an *in-silico* site saturation mutagenesis of the major interface residues within LD3 (A130 – A139 based on PDB 6IWB; **Figure 2a**). Bcl-2 was left untouched so as not to disrupt venetoclax binding. For experimental characterization, we selected five LD3 amino acids displaying a range in ΔΔG: (i) Q132 and A139 as mild targets (mean change in ΔΔG = 0.1 each), (ii) D138 as a highly disruptive target (mean change in ΔΔG = 6.8), and, (iii) a combination of A130 and R134, both medium-range targets (median change in ΔΔG = 3.4 & 1.3, respectively), thereby generating a total of 460 variants, including R134A and D138A, for which we had previously observed increased drug sensitivity^28^ (**Figure S2**). These variants were directly integrated into the corresponding genomic region of LD3 within the anti-HER2 DROP-CAR B3Z cells by Cas9-mediated homology-directed repair (HDR) (**Figure 2b**).

**Figure 2:**
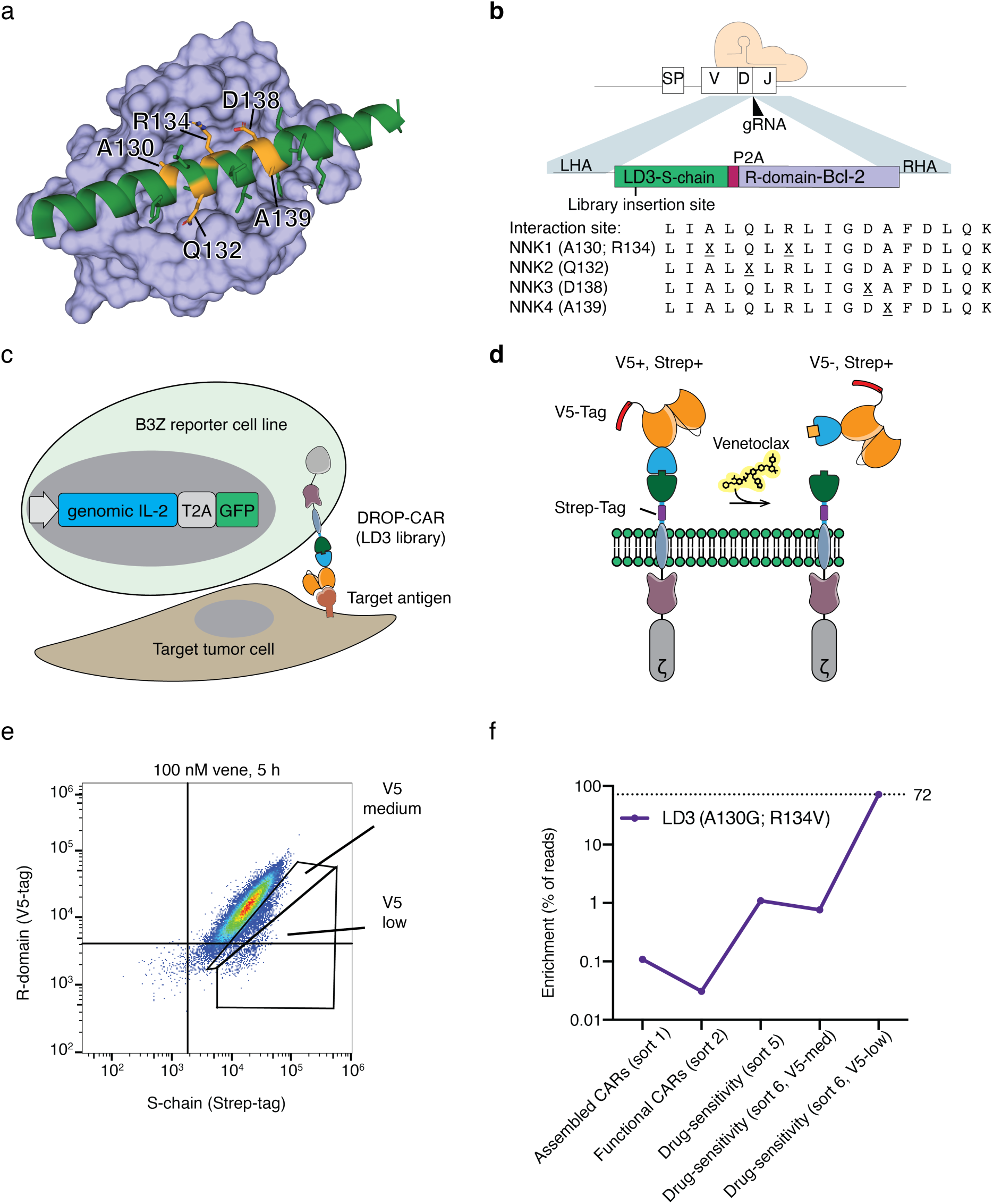
Library design and screening to enrich for LD3 variants enabling DROP-CAR stability coupled with sensitivity to disruption by venetoclax. **a,** Major interaction site between Bcl-2 (cyan) and LD3 (green). Residues chosen for mutagenesis are highlighted in orange. The model is based on PDB 6IWB; only the major interacting helix of LD3 is shown. **b,** Site saturation mutagenesis library for three single interface LD3 variants and one double mutant. The codons for residues marked by X were replaced with the NNK degenerate codon. **c,** Schematic representation of coculture of DROP-CAR B3Z cells with HER2^+^ SKOV3 cells and selection of functional DROP-CARs (GFP^+^ cells). **d,** Illustration of tagged S-chain and R-domain and DROP-CAR disassembly upon venetoclax administration. **e,** Sorting for medium and low levels of V5-tag labeling (from sort 6). **f,** Enrichment of the A130G+R134V LD3 variant after indicated sorts as analyzed by deep sequencing.

In total, we performed six rounds of screening. First, we sorted the library for cell-surface expression of both the S-chain and R-domain (i.e., DROP-CAR cell-surface assembly) by virtue of dual tag labeling (sort 1, **Figure S3**). Subsequently, to remove LD3 variants generating non-functional DROP-CARs, we cocultured the cells from sort 1 with the HER2^+^ ovarian tumor cell line SKOV3 and selected for activated library cells via expression of a GFP reporter gene (genomically integrated downstream of exon 4 of IL-2). This was done twice (sorts 2 & 3)^12^ (**Figure 2c, Figure S3**). We then conducted 3 sequential library screenings for venetoclax sensitivity (sorts 4 - 6). Briefly, for the sensitivity screenings, upon 5 h incubation with 100 nM venetoclax, the cells were labeled with antibodies targeting the Strep-tag (on the S-chain) and the V5-tag (on the R-domain) and we sorted for cells with decreased presence of the R-domain (i.e., the R-domain had been released upon venetoclax incubation) relative to the S-chain (**Figure 2d & e, Figure S4**).

Sequence analysis was performed for: (i) sort 1 to ascertain the diversity of LD3 variants that maintain Bcl-2 binding, (ii) sort 3 to evaluate LD3 variants present in functional DROP-CARs, (iii) sort 5 to evaluate the range of LD3 variants sensitive to venetoclax disruption, and for (iv) sort 6 we sequenced V5-medium and V5-low labeled cells to compare LD3 variants having moderate and high sensitivity to venetoclax disruption. Consistent with our *in silico* analysis, functional CARs enriched in the library screening after sort 3 largely comprised variants of the two sites predicted to be least disruptive to LD3:Bcl-2 complex formation, Q132X and A139X. Few variants were identified for the position predicted to be most disruptive, D138X. A wide range of variants were enriched for the double mutant A130X + R134X, from undetectable to 1.4% (**Table S4**).

In the third and final drug sensitivity screen (sort 6), a single double variant, A130G + R134V dominated, making up just over 70% of the most drug-sensitive population (V5-low). In earlier sorts, this variant was present at 0.11% (sort 1: assembled CARs), 0.03% (sort 3: functional CARs), 1.1% (sort 5: second drug-sensitivity sort), and 0.76% (sort 6: third drug-sensitivity sort, V5-medium) of the population (**Figure 2f, Table S4**). The remarkable enrichment of the A130G+R134V variant of LD3 after the 3 drug-sensitivity screens suggests that the optimal protein interface enabling both DROP-CAR stability and drug sensitivity of the PPI is sequence-limited.

### Selection of the best performing DROP-CAR and affinity measurement of LD3 variants for Bcl-2

We next generated mutant DROP-CAR B3Z cells for the three most enriched variants from the third drug-sensitivity selection (sort 6, V5-low; **Figure S4c,d**), including the LD3 variant A130G+R134V which we named double mutant (dm)LD3, as well as single mutants Q132L and A139R, and evaluated venetoclax-induced receptor disruption. We observed minor disruption of DROP-CARs comprising wild-type LD3, partial disruption for both the Q132L and A139R variants and major disruption for the double mutant A130G+R134V (dmLD3, **Figure S5**). The superiority of the double mutant is consistent with its high enrichment during the library screening. As mentioned, LD3 comprises a human protein scaffold (Apolipoprotein E4; ApoE4) computationally engrafted with critical binding residues of the BH3 domain of Bim^22^. dmLD3 hence generates a human protein having a total of only 12 amino acid difference from wild type (of ApoE4; **Figure S6**).

Finally, we determined the K_D_ of the best LD3 variant, dmLD3, versus the single mutant LD3 (smLD3) variant A139R, with Bcl-2. Recombinant proteins were produced, purified and evaluated for binding affinity by surface plasmon resonance (SPR). We calculated a K_D_ of 82 nM for dmLD3 & Bcl-2, and a K_D_ of 67 nM for smLD3 & Bcl-2 (**Figure S7a & b**), both weaker binding (as expected) in comparison to our previously calculated K_D_ of 0.8 nM for wild-type LD3 & Bcl-2 which is poorly disrupted by venetoclax.

### Characterization of DROP-CAR T cells comprising the optimized dmLD3 variant

Next, we sought to build a prostate specific membrane antigen (PSMA)-targeted DROP-CAR comprising the J591 scFv^30^, human signaling components, and the optimized dmLDR:Bcl-2 off-switch, and to test it for expression and function (**Figure S7c**). We began by engineering Jurkat cells and demonstrated near-total disassembly of the DROP-CAR after 24-hour incubation with 1 µM venetoclax, yielding an IC_50_ of 59 nM (95% confidence interval of 54 to 64 nM; **Figure 3a**). In contrast, we observed no impact of venetoclax on the expression of an equivalent 2G-CAR (**Figure S7d, e**). Our findings are in range with the standard oral dose of 400 mg venetoclax given to patients, which typically achieves a plasma concentration of 2 µM within 6 hours^31, 32^. We further investigated the speed of DROP-CAR disruption. With a venetoclax concentration of 1.25 µM, we observed a receptor disassembly half-life of approximately 2.8 hours (95% confidence interval of 2.6 to 3.1 hours; **Figure 3b**). Given that venetoclax has a terminal half-life of approximately 19-26 hours in humans^31, 32^, this response rate suggests compatibility with a treatment regimen of low-dose venetoclax that can be adjusted to balance DROP-CAR T-cell function and safety.

**Figure 3:**
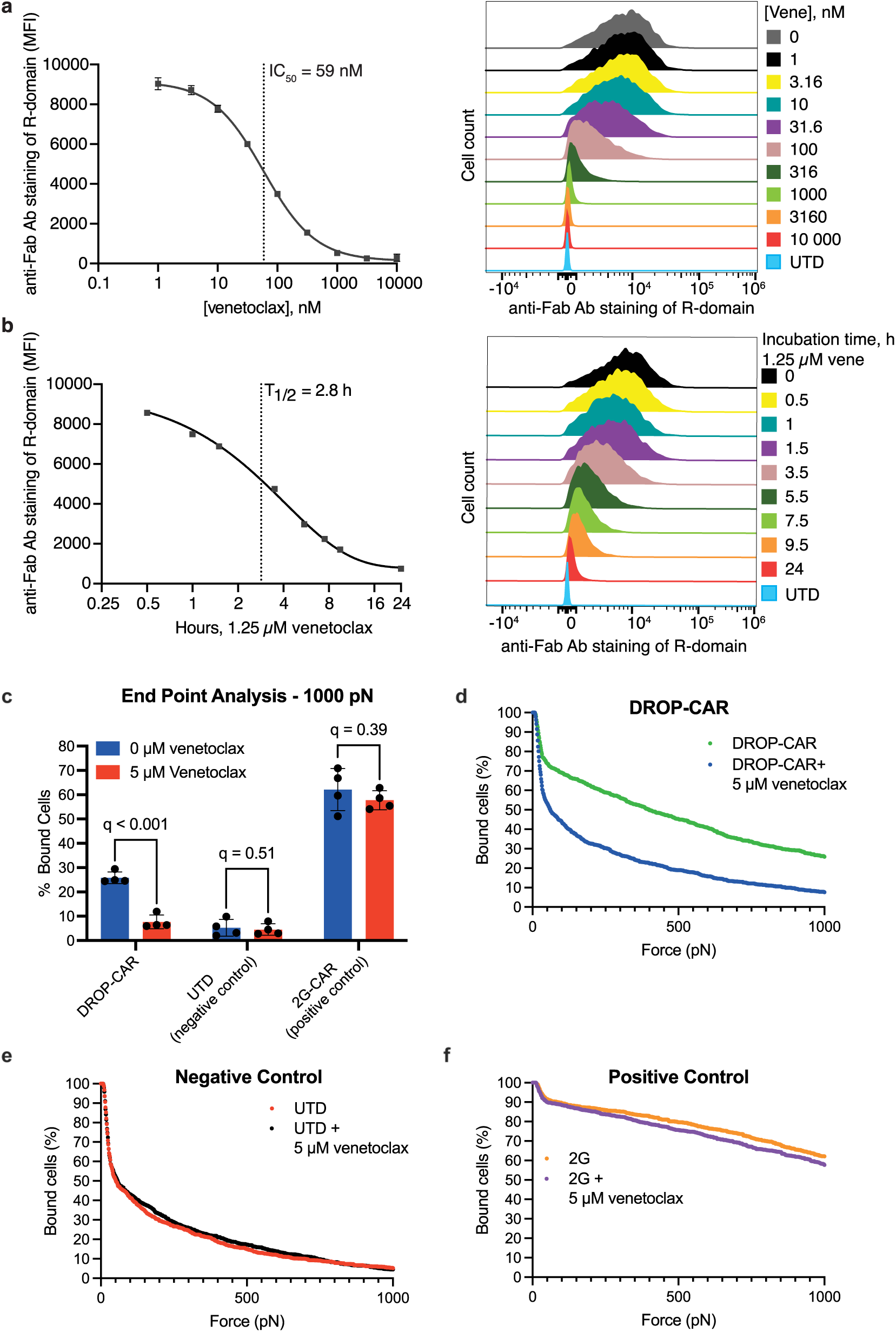
Characterization of optimized DROP-CAR and proof-of-principle for dual DROP-CAR cells. **a,** Dose-response curve of optimized DROP-CAR (comprising dmLD3) Jurkat cells in response to venetoclax. **b,** Time-course experiment of optimized DROP-CAR Jurkat cells at indicated time points after incubation with 1.25 µM venetoclax. Transduction efficiency for the DROP CAR was 100%. **c,** Percentage of Jurkat cells that remain bound after 2.5 min of a linear acoustic force ramp from 0 to 1000 pN. Partly detached Jurkat cells (hinge cells) are counted as detached. Values are the mean ± s.d. of n = 4 measurements on separate chips. **d,e,f** Percent of cells bound for the DROP-CAR Jurkat cells, untransfected cells (UTD), and 2G-CAR Jurkat cells over 2.5 min of a linear acoustic force ramp from 0 to 1000 pN. Transduction efficiency for the DROP CAR was 59% and for the 2G CAR 99%. Partly detached cells (hinge cells) are counted as detached. Values are n = 4 measurements on separate chips.

### Cell avidity measurements to test for cell-cell contact regulation

We postulated that abrogation of DROP-CAR T-cell activity upon venetoclax administration is due to release of the R-domain and breakage of cell-cell contact. Hence, we next measured avidity between anti-PSMA DROP-CAR Jurkat cells and target PC3-PIP -/+ venetoclax cells using a LUMICKS zMovi device. For the assay, T-cells were added onto a monolayer of cancer cells, allowed 5 min to bind, and then detachment tracked by microscopy over 2.5 min while applying a force ramp from 0 to 1000 pN. We observed a venetoclax-induced reduction in the number of bound DROP-CAR-but not 2G-CAR-Jurkat cells (**Figure 3c-f**). If partly detached cells were counted as dissociated, in the presence of venetoclax both UTD and DROP-CAR Jurkat cells approached less than 10% bound while 2G-CAR Jurkat cells were 60% bound. DROP-CAR Jurkat cells remained about 30% bound in the absence of venetoclax. In contrast, when partly detached cells were counted as bound, at 1000 pN in the presence of venetoclax UTD cells were approximately 25% bound, DROP-CAR Jurkat cells 30%, and 2G-CAR Jurkat cells just under 90%. In the absence of venetoclax, DROP-CAR Jurkat cells remained 64% bound (**Figure S8**). These results are consistent with the transduction efficiency for this experiment (DROP-CAR: 59%, 2G-CAR: 99%) and confirm that cell-cell interactions can be regulated for DROP-CAR T cells.

### Generation of independently controlled dual DROP-CAR T cells

A variety of approaches, including dual CARs and Tandem CARs which engage more than one antigen^33^, and adapter/universal CARs for transiently targeting antigen(s) of choice^24, 34^, have been developed to overcome antigen escape which is an important barrier to CAR therapy^5^. Hence, we next sought to test proof-of-concept for DROP-CARs targeting two different antigens. Briefly, we built a lentiviral construct encoding the S-chain comprising dmLD3 followed by two different R-domains. The first R-domain targeted the B-cell lineage antigen CD19 (FMC63 scFv)^35^ and was fused to Bcl-xL (A-1155463^22^ responsive), and the second one targeted PSMA (J591 scFv)^30^ and was fused to Bcl-2 (venetoclax responsive) (**Figure S9a & b**). We tagged Bcl-xL with V5 and Bcl-2 with HA to allow us to track the disruption/loss of each R-domain upon small molecule administration. A-1155463 caused almost complete release of Bcl-xL while venetoclax caused only partial release (**Figure S9c**) consistent with its moderate inhibition of Bcl-xL (Ki = 48 µM)^36^. Venetoclax strongly released Bcl-2 but A-1155463 almost not at all (**Figure S9d**). While this dual system is not clinically relevant because A-1155463 is not approved, it demonstrates that it is feasible to differentially modulate the expression of two different DROP-CARs with 2 small molecules.

### Extending the DROP-mechanism to other synthetic receptors

We next sought to evaluate if the DROP-mechanism could be extended to other synthetic receptor-types and thus built it into the generalized extracellular molecule sensor (GEMS) cytokine receptor platform and tested it in HEK293T cells. Briefly, GEMS comprises the extracellular domain of the erythropoietin receptor (EpoR) fused to the intracellular domain of IL6R-beta. To enable inducible dimerization, domains that respond to specific stimuli (such as nanobodies that dimerize in response to caffeine) are fused to the N-terminus of the EpoR domain^37, 38^

We introduced the dmLD3:Bcl-2 module between the EpoR and caffeine nanobodies with the aim of creating a system that turns on in response to caffeine (dimerization of the nanobodies) and turns off in response to venetoclax (by causing nanobody release; **Figure S10a**). The system worked with the intended behavior and we observed enhanced shut-off for dmLD3 versus wild-type LD3 (**Figure S10b**). This confirms that dmLD3 forms complexes that are stable enough to yield functional cytokine receptors as well as CARs, and can also be disrupted more efficiently than wild-type. We subsequently confirmed that Bcl-xL and A-1155463 are also functional in this context (**Figure S10c**), showing that the DROP-mechanism can be extended other synthetic receptor platforms and could be useful for building systems with elaborate logics. Notably, this type of ON/OFF logic for two soluble molecules can be referred to as A AND (NOT B) or A NIMPLY B in a synthetic biology context^39^ (**Figure S10d**). Such mechanisms may be useful to turn on receptor activity or gene expression in response to environmental cues or inducer molecules, but only if a second condition is not present.

### In vitro and in vivo function of human DROP-CAR T cells as compared to 2G-CAR T cells

Lastly, we sought to compare human DROP-CAR-versus equivalent 2G-CAR T cells^40^. For the DROP-CAR we achieved 40-60% lentiviral transduction efficiency in CD4^+^ T cells and 20-40% in CD8^+^ T cells, versus about 80% transduction efficiency for the 2G-CAR in both CD4^+^ and CD8^+^ T cells (**Figure S11a**). These values are in line with the general observation that transduction efficiency decreases with the size and complexity of the CAR^33, 41^ but are well above typical thresholds for clinical trials which can be set below 10% as transduced cells are expected to expand in the presence of antigen^42^. A higher MFI was observed for the 2G CAR indicating that it is expressed at higher levels per cell than the DROP-CAR (**Figure S11b**), but lower cell-surface density may in fact be beneficial in a clinical context^43^. We further evaluated memory phenotype and expansion at 11 days post-transduction but noted no significant differences amongst the CAR T cells and untransduced (UTD) T cells (**Figure S11c-e**).

In terms of in vitro function, both 2G- and DROP-CAR T cells produced IFN-γ upon co-culture with target PSMA^+^ PC3-PIP tumor cells (**Figure 4a**) albeit at lower levels for the latter (at 24 hours), corresponding to lower expression levels of the DROP-CAR (**Figure S11b**). Importantly, co-culture in the presence of 1.25 µM venetoclax abrogated IFN-γ production by DROP-CAR T cells but did not hinder the 2G ones (**Figure 4a**). We subsequently set-up IncuCyte real-time cytotoxicity assays and demonstrated robust and equivalent killing of PC3-PIP tumor cells by 2G- and DROP-CAR T cells, and a dose-dependent decrease in cytotoxicity by DROP-CAR-but not 2G-CAR T cells in the presence of venetoclax (**Figure 4b,c**). Co-culture with UTD T cells +/- venetoclax had no impact on the target tumor cells (**Figure 4b,c**).

**Figure 4:**
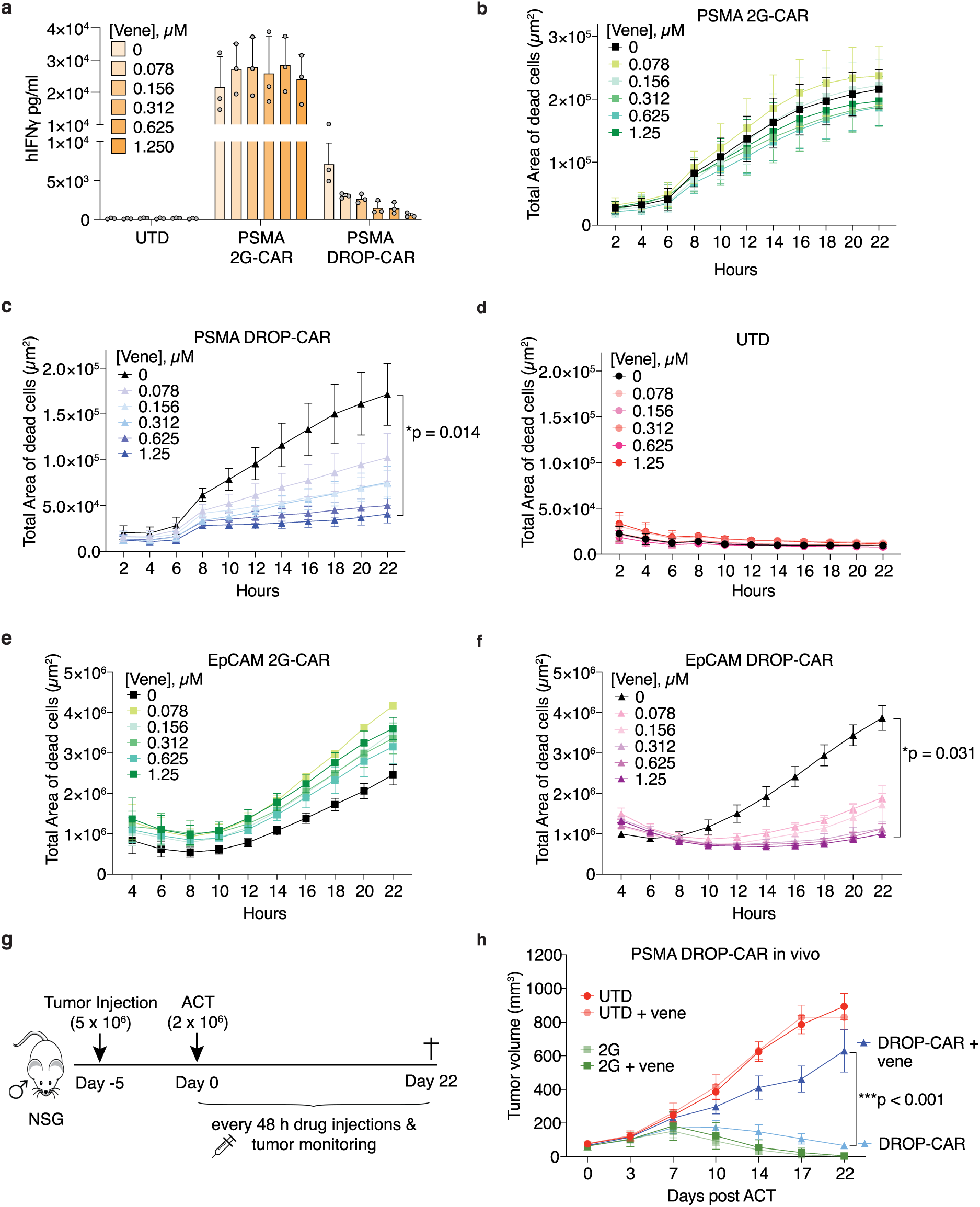
In vitro and in vivo characterization of human DROP-CAR T cells. **a,** IFN-γ production by anti-PSMA CAR T cells upon 24 hours coculture with PSMA^+^ PC3-PIP cells +/- venetoclax (ven). **b,** IncuCyte cytotoxicity assay at an effector to target ratio of 2:1 to evaluate PC3-PIP tumor cell killing by anti-PSMA 2G-CAR T cells, **c**, DROP-CAR T cells, or, **d,** UTD T cells in the presence of up to 1.25 μM venetoclax. **e,** IncuCyte cytotoxicity assay to evaluate (EpCAM^+^) PC3-PIP tumor cell killing by anti-EpCAM 2G-CAR T cells or **c**, DROP-CAR T cells in the presence of up to 1.25 μM venetoclax. Values are the mean ± s.e.m. of n = 3 (**a,e,f**), n = 8 (**b,c,d**) human donors. **g,** Schematic of in vivo adoptive T cell transfer (ACT) study (n = 7 mice/group). **h,** Tumor control curves over 22 days for the ACT study. Before all functional assays, the engineered T cells were rested and adjusted by mixing them with UTD cells for equivalent transgene expression levels. A mix of 50% CD4^+^ T cells and 50% CD8^+^ T cells was used in all functional experiments.

We next built lentiviral vectors encoding 2G- and DROP-CARs targeting epithelial cellular adhesion molecule (EpCAM) with scFv C215^44^, also comprising a CD8α hinge, the TM and endodomain derived from CD28, and the endodomain of CD3ζ. We achieved similar transduction efficiencies as for the anti-PSMA CARs (**Figure S11f**) and the CAR T cells were able to robustly kill PC3-PIP cells, which are naturally EpCAM^+^, with only the DROP-CAR T cells showing sensitivity to venetoclax (**Figure 4e,f, Figure S11g**).

Encouraged by the high cytolytic capacity of DROP-CAR T cells, we next sought to compare subcutaneous PC3-PIP tumor control in NSG mice by anti-PSMA 2G-versus DROP-CAR T cells +/- venetoclax (**Figure 4g**). We observed that both 2G- and DROP-CAR T cells can efficiently control PC3-PIP tumors and that venetoclax administration (2.5 mg/kg every 48 hours) abrogated control by the DROP-CAR T cells but had no impact on the 2G-CAR T cells in vivo (**Figure 4h**). Importantly, we showed that the suppression is reversible as upon venetoclax withdrawal DROP-CAR T cells regained tumor control, even at an advanced stage (**Figure S12a**). In an independent in vivo study, we observed that 2.5 mg/kg of venetoclax did not fully abrogate DROP-CAR T cell activity, but that increasing the dose to 5 mg/ml restrained their function (i.e., the tumors escaped) (**Figure S12b**). Remarkably, upon venetoclax withdrawal at day 42, the tumors began regressing in the treated mice, indicating that the DROP-CAR T cells were still present and had regained function (**Figure S12c**).

## Discussion

Here, by structure guided design and mammalian cell library screening, we have developed an optimized off-switch comprising human protein components that form a stable PPI that can be efficiently disrupted by the clinically approved small molecule venetoclax. We further demonstrated remote-control of human DROP-CAR (off-switch) T cells in tumor-bearing mice. At the outset of this study, we envisioned a DROP-CAR design made up of a single transmembrane chain encoding intracellular signaling components (the S-chain) along with a soluble domain including an anti-tumor antigen scFv (the R-domain) that associate via a disruptable PPI. Importantly, we demonstrated that venetoclax administration prevents cell-cell contact between the DROP-CAR T cell and target tumor cell and reversibly abrogates T-cell function. Our design is akin to a modular universal CAR except that for our strategy the tumor-antigen binding adaptor protein (i.e., the R-domain) is produced by the engineered T cells themselves. In contrast, universal CARs rely upon frequent injections of soluble adaptor protein dosed at mg/kg levels^24^ which can increase the risk of off-target effects. In addition, there are the added expenses and logistics of producing and administering the adaptor protein, along with potential barriers to deep tumor penetration and encounter of engineered T cells^5^.

For our PPI, we began with truncated Bcl-2 which is structurally similar to Bcl-xL and also binds to our human designer protein LD3 with high-affinity^28^. However, LD3 could not be readily dissociated from Bcl-2 by venetoclax. Hence, we built a mammalian cell-based library of DROP-CARs with amino acid replacements at selected sites in LD3 predicted to be minimally to highly disruptive to Bcl-2 binding. The screening was performed in a logical manner. First, we selected for cells expressing both chains at their surface, then for ones expressing a functional DROP-CAR, and finally we enriched for stable DROP-CAR variants that are also sensitive to disruption by venetoclax. Our library design and screening strategy might readily be applied to the optimization of other disruptable PPIs.

Remarkably, of the nearly 500 LD3 variants built into the library, only one was highly enriched, dmLD3 (A130G+R134V) which could maintain stability of the PPI while increasing sensitivity of the interface to venetoclax disruption. Another variant (A139R) that emerged in the screening was in a similar range of binding affinity as dmLD3 (A139R: 67 nM, dmLD3: 82 nM) for Bcl-2, but was less sensitive to venetoclax. This observation suggests that additional factors for dmLD3, such as potentially higher flexibility of the binding site, increased hydrophobicity of a potential drug-entry point, or/and other effects, may contribute to more sensitive switching. Our computational ΔΔG predictions did not reveal any unique features of these mutations, underscoring the limitations of rigid body docking for this application and highlighting the need for the developed screening approach for selecting the best-performing DROP-CAR. In total, dmLD3 includes the engraftment of 12 amino acids (i.e., 10 critical binding residues from the BH3 domain of Bim^22^ and 2 selected in the screening here) onto the scaffold protein ApoE. We propose that risk of immunogenicity of this fusion human protein is low, especially considering that the 12 amino acids are located within a protein interface.

We successfully engineered human T cells to express DROP-CARs targeting PSMA as well as EpCAM and demonstrated robust in vitro function against target tumor cells, but not in the presence of venetoclax which disrupts the PPI (to release the R-domain) and can bind Bcl-2 with picomolar affinity^45^. Presumably the DROP-CAR assembles within the endoplasmic reticulum and remains sufficiently stable at the T-cell surface until renewed by natural protein turnover. Others have estimated a CAR turn-over half-life of 6h^46^. Importantly, we also showed subcutaneous solid tumor control and that DROP-CAR T-cell function could be reversibly blocked upon administration and withdrawal of venetoclax. In vitro, the optimized DROP-CAR design could be disrupted with a half-life of about 3 hours and an IC_50_ of 59 nM. These values are well below the typical plasma concentrations of about 2 µM venetoclax 6 hours after the standard oral dose of 400 mg and the 19-26 h terminal half-life^31, 32^. These characteristics and the observed dose dependence on abrogation of target-cell killing suggest compatibility with a potential daily or bi-daily regimen of low-dose venetoclax optimized for the highest on-tumor and tolerable off-tumor toxicity. Futures studies should include testing the ability to abrogate DROP-CAR T-cell exhaustion by transient resting^6^ as well as to mitigate toxicity in vivo.

Notably, we demonstrated the modularity of the switch by generating caffeine-induced engineered cytokine receptors based on the GEMS platform^37, 38^. Such a response pattern to two small molecules (i.e., caffeine and venetoclax) serves as proof-of-concept for AND-NOT logic, which is one of the fundamental logic gates for circuit design in synthetic biology^39^. Moreover, it demonstrates that the DROP mechanism can be used to add additional layers of control to other receptor systems beyond CARs. This feature could be of interest for various controllable intra- and extracellular receptor systems including inducible zinc finger transcription factors^16^, the recently humanized latest generation of SynNotch receptors^47^ and other synthetic receptors^48,49^. Additionally, we showed that the off-switches can be extended to display more than one scFv. Current tandem receptor designs that recognize two tumor antigens are highly promising for mitigating antigen escape^50^ but reliable cancer cell surface markers are rare and the simultaneous targeting of multiple markers might exacerbate on-target, off-tumor toxicities. scFv-specific control options by dual-DROP-CARs could therefore offer a way to mitigate risks associated with CAR T-cell therapies aimed at minimizing antigen escape. Taken together, our work makes important inroads towards optimizing the function and safety of cellular gene-therapies for cancer and beyond.

## Methods

### *In silico* LD3 interface site saturation mutagenesis

The Rosetta modeling suite was used to compute the ΔΔG for the LD3:Bcl-2 complex (PDB 6IWB) upon mutating each of the 10 major interface residues to all possible 20 amino acids. The structure was relaxed with the ’FastRelax’ mover to calculate the initial ΔΔG. Each interface residue of interest was mutated by side chain repacking and energy minimization using ’PackRotamersMover’ and ’MinMover’ respectively. The difference of ΔΔG between the mutated and initial state was then calculated. According to our computational workflow, ΔG refers to the change in Gibbs free energy upon protein folding, while ΔΔG refers to the change upon protein complex formation, both expressed in Rosetta energy units. We compare the ΔΔG values resulting from *in silico* site saturation mutagenesis to the ΔΔG value from the wild-type complex (change in ΔΔG = ΔΔG_mutant_ - ΔΔG_WT_). A positive change in ΔΔG indicates a weakened protein interface, predicting a reduction in the binding affinity of the mutated complex compared to the wild-type.We repeated the same method 3 times to calculate the mean and standard deviation of the results. Based on the results and the structure, we did not mutate L131 and L135 as they don’t point toward the interface and might contribute to stabilizing the LD3 core. We discarded G137 as almost every mutant results in clashes with Bcl-2, which likely is prohibitive of binding. From the remaining sites, we experimentally tested two mildly disruptive mutants (Q132, A139), one strongly disruptive mutant (D138), and two mutants that showed a wide spread of ΔΔG (130A, 134R). We tested 130A and 134R mutations in combination to maximize the tested range of different affinities.

### Compound preparation and storage

Venetoclax (>99.9%, Chemietek, CT-A199) and A1155463 (99.5%, Chemietek, CT-A115) were prepared in DMSO (Neofroxx, 1264) as 10 mM stocks and aliquots were stored at -20 °C. Caffeine was prepared in Milli-Q water as 100 mM stocks and aliquots were stored at -20 °C.

### Protein structure visualizations

The models in Figures 1 and 2a were generated with PyMol^40^. Figure 1a was assembled from AlphaFold2 ^51^ models of individual receptor domains for visualization purposes and is not an accurate portrayal of a real protein structure.

### Cell culture

The prostate carcinoma cell line PC3-PIP and Jurkat T cell line were cultured in RPMI-1640 (Gibco, 72400047). B3Z cells were cultured in IMDM (Gibco, 31980030). HEK293T cells were cultured in DMEM (Gibco, 10566016) for experiments and in RPMI-1640 for virus production. SKOV3 cells were cultured in DMEM/F-12 (Gibco, 31331028). All medium was supplemented with heat-inactivated 10% FBS (Gibco, 26140079) and Pen/Strep (Gibco, 10378016). All cells were cultured at 37°C and 5% CO_2_ in a humidified incubator.

### HEK293T cell experiments and reporter assay

For the HEK293T cell experiments and reporter gene assay (in **Figure 4**), 16,000 cells per well were seeded with 100 µL of complete DMEM medium in the inner 60 wells of a 96-well plate, 24 h before transfection. The DNA mix for 12 wells consisted of 600 ng receptor plasmid (S193), 600 ng secreted LD3-nanobody fusion (pLS147 or pLS912), 200 ng STAT3 expression plasmid (pLS392), 300 ng reporter plasmid for STAT3 dependent nanoluciferase secretion, mixed with 660 µL DMEM without additives and 8250 ng polyethyleneimine (PEI; Polysciences Inc., 24765-1). All plasmids are described in **Table S1**. The transfection mixture was vortexed, incubated for 20 minutes, and 50 µL/well was added to the cells. These numbers correspond to ∼130 ng DNA and 625 ng PEI per well. After 16 hours, the medium with the transfection mix was replaced with 100 µL per well of fresh complete medium containing the indicated drugs. After 2 h, supernatant samples were collected for quantification of the secreted reporter protein nanoluciferase. For the assay, 5 µL of cell supernatant was mixed with 5 µL of a nanoglo substrate:buffer solution (Promega, N1110) at a 1:50 ratio in a black 384-well plate and analyzed using a multiplate reader. For a more detailed protocol, please refer to ^38^.

### Protein expression and purification

Both dmLD3 (A130G; R134V) and smLD3 (A139R) with a C-terminal His_6_-tag were produced using the Expi293TM expression system (Thermo Fisher Scientific, A14635). Six days post-transfection, the supernatant was harvested, filtered, and the protein was purified using Ni-NTA affinity columns on an ÄKTA pure system (GE healthcare), followed by size exclusion chromatography in PBS. The purified proteins were then concentrated, aliquoted, and stored at -80 °C for subsequent experiments. The Bcl-2 protein for SPR experiments was produced previously ^52^.

### SPR assay for protein-protein binding affinities

To determine the binding affinities between the LD3 variants and Bcl-2 proteins, surface plasmon resonance (SPR) measurements were conducted using a Biacore 8 K (GE Healthcare). The Bcl-2 protein was immobilized on a CM5 chip (GE Life Science) at 5 µg/ml for 140 seconds of contact time in pH 4.5 sodium acetate solutions. Serial dilutions of the mutated LD3 proteins were prepared in HBS-EP buffer (10 mM HEPES, 150 mM NaCl, 3 mM EDTA, and 0.005% Surfactant P20; GE Life Science) and flown over the immobilized chips. The binding affinities (dissociation constant; K_D_) were determined using a steady binding model or equilibrium model with the Biacore 8 K evaluation software.

### Plasmid construction

All plasmids are described in **Table S1** and all relevant protein sequences are listed in **Table S2**. DNA for cloning was ordered from Twist Bioscience, either as gene fragments or as readily cloned vectors. The scFv-bcl-xL insert in the dual-Drop-CAR was purchased from GenScript. Gene fragments were ordered with flanking regions overlapping with the target vector and inserted by Gibson assembly. The initial DROP-CAR DNA was cloned into a pUC57 vector containing the homology arms for homology-directed repair in the B3Z cells. The DROP-CARs for Jurkat and primary T cell experiments were cloned into a third-generation self-inactivating lentiviral expression vector, pELNS, with expression driven by the strong elongation factor-1α (EF-1α) promoter. Recombinant LD3 variants were expressed from a pHLSec vector with a strong CAG promoter. GEMS and secreted LD3-nanobody fusion proteins were expressed from plasmids with a weak SV40 promoter.

### Lentivirus production

24 hours before transfection, 10 million HEK-293 T cells were seeded in 16.5 ml of complete DMEM in a T-150 tissue culture flask. Plasmid DNA was purified using the Endo-free Maxiprep kit (Qiagen, 12362). For transfection, a mixture of 7 µg pVSV-G (VSV glycoprotein expression plasmid), 18 µg R874 (Rev and Gag/Pol expression plasmid), and 15 µg pELNS-based vector was combined with 180 µl Turbofect (Thermo Fisher Scientific AG, R0533) and 3 ml Optimem medium (Gibco, 31985062). After 48 hours of incubation, the viral supernatant was collected, and viral particles were concentrated by ultracentrifugation for 2 hours at 24,000 g, then resuspended in 400 µl of complete RPMI-1640 medium and snap-frozen on dry ice.

### Jurkat cell Transduction

Jurkat cells were seeded at a density of 1 million cells/ml in 48-well plates with 500 µl per well. For each transduction, 50 µl of viral supernatant was added to the wells. Following 24 hours of incubation at 37°C, the cell medium was replaced, and the cells were incubated for an additional 72 hours at 37°C.

### Primary human T-cell transduction and culture

Primary human T cells were extracted from the peripheral blood mononuclear cells (PBMCs) of healthy donors (HDs). PBMCs were separated using Lymphoprep (Axonlab, 12053231) and CD4+ and CD8+ T cells were isolated using a magnetic bead-based negative selection kit (easySEP, Stem Cell technology, 17951) according to the manufacturer’s instructions. The isolated T cells were cultured in complete RPMI and stimulated with Dynabeads™ Human T-Activator CD3/CD28 for T Cell Expansion and Activation (Gibco, 11132D) at a 1:2 ratio of T cells to beads. T cells were transduced (0.25 million CD4+ and CD8+ cells for in vitro experiments and 3 million for in vivo) with lentiviral particles 18-22 hours after activation. Human recombinant IL-2 (PeproTech, 200-02) was added every other day at a concentration of 50 IU/ml until day 5 post-stimulation. On day 5, the beads were removed and IL-2 was replaced with IL-7 and IL-15 (Miltenyi Biotec GmbH, 130-095-367 and 130-096-567) at 10 ng/ml each. The cells were maintained at a density of 0.5-1 million cells/ml for expansion until day 11 or 12. Before functional assays, engineered T cells were rested and mixed with UTD cells to standardize transgene expression levels. Unless stated otherwise, a mix of 50% CD4+ T cells and 50% CD8+ T cells was used in all functional experiments.

### Cytokine production assays

For cytokine release assays, an effector to target ratio of 1:1 was used. Briefly, 50,000 T cells per well were co-cultured with an equal number of target cells in 96-well round-bottom plates in a final volume of 200 µl of complete RPMI medium. After 24 hours, supernatant samples were collected to measure IFN-γ and IL-2 using commercial enzyme-linked immunosorbent assay (ELISA) kits according to the manufacturer’s instructions (BioLegend, 430801 and 431801).

### Cytotoxicity assays

Cytotoxicity assays were conducted using an IncuCyte Instrument at a 2:1 effector to target ratio (50% CD4^+^ and 50% CD8^+^ T cells) in complete RPMI medium containing Cytotox Red reagent (final concentration of 125 nM; Essen Bioscience, 4632) without exogenous cytokines. Target cells (12,500) were seeded in flat-bottom 96-well plates, and after 4 hours of incubation, 25,000 washed and rested T cells (no cytokine addition for 48 hours) were added per well in a final volume of 200 µL. Images were taken every 2 hours, and the total red area per well was calculated using IncuCyte ZOOM software. Cytotoxicity is reported as the total area of dead cells as measured by Cytotox Red reagent uptake (red area per µm²). Background fluorescence at time 0 was subtracted from all subsequent time points. Data are presented as the mean of total red area for different healthy donor cells ± s.e.m.

### Mice and In Vivo experiments

NOD SCIDγ (NSG) mice were bred and housed in a pathogen-free facility at the Epalinges Campus of the University of Lausanne. All experiments followed Swiss Federal Veterinary Office guidelines and were approved by the Cantonal Veterinary Office. Five mice were housed per cage, provided an enriched environment and unrestricted access to food and water. Mice were monitored at least every two days for signs of distress and were euthanized using carbon dioxide overdose at the endpoint. To evaluate tumor control by DROP-CAR-T cells, 8-12 week-old NSG males were subcutaneously injected with 5 million PC3-PIP tumor cells. Groups of 7-9 mice (as indicated in the figure legends) were formed once tumors became palpable (day 5) to ensure similar mean tumor volume and standard deviation. The mice were treated with peritumoral injection of 2 million T cells (UTD, 2G-CAR, or DROP-CAR-T cells). Two hours after T-cell transfer the mice received peritumoral injections of either 2.5 mg/kg venetoclax or vehicle control (DMSO) every two days until the endpoint or as indicated to evaluate the regain of function of DROP-CAR T cells. Tumor volumes were measured every two days using the formula V = 1/2 (length × width²), where length is the greatest longitudinal diameter and width is the greatest transverse diameter, determined via caliper measurement.

### Genome editing

All sequences related to genome editing are listed in **Table S3**. To generate the initial DROP-CAR in stably Cas9 expressing B3Z cells, purified PCR product of the gene cassette with ∼700 bp flanking homology arms left and right of the genomic cut site was used for the transfection as the template for homology-directed repair (HDR). For library generation, we used short 125-base single-stranded oligodeoxynucleotides (ssODNs; 500 pmol ultramer; IDT) with about 60 bases left and right as homology regions and phosphorothioate bonds at each end as the template. Electroporations were performed with the SF Lonza kit on a Lonza 4D-Nucleofector by resuspending 5 × 10^4^ B3Z cells in SF buffer in a total volume of 100 µL and running program CA-138 in nucleocuvettes, followed by the addition of 600 µL warm complete RPMI medium. To generate the gRNA, we heated and assembled 2.25 µL of 200 µM tracRNA and 2.25 µL of 200µM crRNA, and transfected 4 µL of the mixture together with 5 µg of template DNA. Analysis or sorting was performed after a minimum of four days. We confirmed successful genomic editing by extracting genomic DNA from at least 10^4^ harvested cells using the QuickExtract protocol (Lucigen, QE09050). Correct HDR template integration was validated through Sanger sequencing of the PCR amplification of the target locus with at least one primer annealing outside the integration site.

### DROP-CAR library generation

A gRNA (**Table S3**) targeting the LD3 region of the initial DROP-CAR was used to introduce a frameshift deletion that removed CAR expression. After sequencing the genomic region of single clones without CAR expression, we selected three clones with small deletions and confirmed the function of the GFP reporter from the genomic IL-2 site by PMA/ionomycin stimulation. We used 4 gRNAs to target the LD3 site in these mutants and repaired this deletion with 125 bp ssODNs as HDR templates containing one or two degenerate NNK codons that encode all 20 amino acids, but only a single stop codon (**Table S3**). The best clone-gRNA-ssODN combination resulted in 15% frameshift repair and 5% HER2 binding recovery.

### Flow cytometry of B3Z cells

DROP-CAR expression in B3Z cells was evaluated via labeling with a 1:200 dilution of biotinylated anti-Strep tag antibody (GenScript, A01737) binding the TMD-containing chain of the DROP-CAR, followed by a 1:400 dilution of Brilliant Violet 421-Streptavidin conjugate (Biolegend, 405225). To select for DROP-CAR assembly, we used a 1:20 dilution of an anti-V5 antibody (Invitrogen, 12-6796-42) that binds to the soluble R-domain (that is released upon venetoclax incubation). Functional DROP-CARs were selected based on inducible GFP expression from the genomic IL-2 site and by HER2 binding (2.5 µg/mL soluble HER2 antigen (Merck) and subsequent 1:200 APC-labeled anti-HER2 antibody (Biolegend)). For this assay, DROP-CAR-library cells were co-cultured in a 1:1 ratio with 3 × 10^6^ HER2-expressing SKOV3 cells in complete IMDM medium for 16 h. All cells were then collected, washed, and sorted for GFP positivity. Cells were allowed to recover for 3 to 5 days before the next selection step. After indicated selection rounds, we amplified the 301 bp LD3-containing region within the CAR gene and sent it for deep sequencing (GENEWIZ). Only sequences with a complete LD3 region harboring a single amino acid substitution were included in the subsequent analysis.

### Dose-response, time course experiments, and flow cytometry of CAR-engineered Jurkat T-cells

Jurkat T-cells (1 × 10^4^ cells/well) were seeded into the inner 60 wells of round-bottom 96-well plates, 24 hours before analysis. For the dose-response experiment and for the 24-hour timepoint of the time course, 50 µL of complete RPMI containing the appropriate doses of venetoclax was added immediately. In the time-course experiment, on the following day, 50 µL of complete RPMI with 3.75 µM venetoclax (yielding a final concentration of 1.25 µM) was added at the specified time points before analysis. CAR assembly was evaluated by labeling with an anti-Fab antibody (1:100, Jackson ImmunoResearch, 115-606-072), which binds to the soluble scFv portion of the receptor. The mean fluorescence intensity (MFI) shift of this population was used to quantify receptor disassembly in response to venetoclax. The V5-Bcl-xL-scFv_CD19_ domain for the dual-DROP-CAR was labeled with an anti-V5 antibody (1:100, Invitrogen, 12-6796-42) to determine receptor disassembly in response to A1155463. The HA-Bcl-2 -scFv_PZ1_ domain for the dual-DROP-CAR was labeled with an anti-HA antibody (1:100, Miltenyi Biotec, 130-120-791) and an anti-mouse secondary antibody (1:100, Invitrogen, 12401082).

### Cell binding force assay

PC3-PIP target cells were harvested using Accutase (PAN-Biotech, Aidenbach, Germany, Cat. No. P10-21200) and seeded on poly-L-lysine (Sigma-Aldrich, Cat. No. P4832, St. Louis, USA) coated z-Movi microfluidic chips (Lumicks, Amsterdam, Netherlands) at 3.5 x 10^7^ cells per ml. Chips with confluent monolayers were incubated for at least 2.5 h at 37 °C prior to measurement of binding force on the z-Movi Cell Avidity Analyzer (Lumicks). Viability of effector cells was assessed by Vi-CELL BLU cell viability analyzer (Beckman Coulter, Brea, USA). Per experimental condition, 1 x 10^6^ effector cells were labelled with a 1 μM Cell Trace far-red reagent (Thermo Fisher Scientific, Waltham, USA, Cat. No. C34564) for 15 min at 37 °C, 5% CO_2_. Effector cells were resuspended at 1 x 10^7^ cells/ml in RPMI-1640 (1X) + GlutaMAX (Thermo Fisher Scientific, Cat. No. 61870-010) supplemented with 10 % FCS (PAN-Biotech, Cat. No. P40-37500), 1 % Penicillin/Streptavidin (PAN-Biotech, Cat. No. P06-07100), and 1 % Hepes (Sigma-Aldrich, Cat. No. H0887).

Binding force was measured and analyzed using Oceon software (V1.5.4) according to manufacturer recommendations. Effector cells were serially flushed into the chips. During a coincubation time of 5 min, effector cells were allowed to form contacts with target cells. Next a force ramp ranging from 0 to 1000 pN was applied over 2.5 min and effector cell detachment was monitored by fluorescent imaging with the z-Movi device. Binding force was determined on single cell level by correlation of the detachment time point with force applied. Runs with <100 or >800 effector cells detected in the measured field of view were excluded A maximum of 8 runs was performed per chip and monolayer viability was assessed by Trypan-blue staining after the last run. Chips with severe viability loss or monolayer cell detachment during the course of the experiment were excluded. During the analysis, effector cells stuck on glass surface or covered by force nodes were automatically deselected, and cells which were moved by force, but did not entirely reach the force nodes, were automatically defined as hinge cells by the Oceon software. Hinge cells were treated as detached cells. Automatic analysis and cell deselection were reviewed but not modified by the experimenter in order to enable reproducibility. In case of severe errors in the automated analysis, chips were excluded from the analysis.

### Flow cytometry of primary T-cells

To evaluate DROP-CAR expression on primary cells, transduced cells were stained with an anti-Fab antibody (Jackson ImmunoResearch, 115-606-072). Cell viability was assessed using a near-infrared fluorescent reactive dye (APC-Cy-7, Invitrogen, Life Technologies, L34976A). For phenotypic memory analysis, the following monoclonal antibodies (mAbs) were utilized: BV711 mouse anti-human CD3 (BD Bioscience, 563725), BV605 mouse anti-human CD4 (Biolegend, 317438), APC-labeled anti-human CD8 (Biolegend, 344722), PE-Texas red-labeled mouse anti-human CD45RA (Beckman Coulter, B49193), and BV421 mouse anti-human CCR7 (Biolegend, 353207). The memory phenotype was determined by first gating the CD3+ population, then separating the CD4+ and CD8+ subsets. These subsets were subsequently analyzed for CD45RA and CCR7 expression to ascertain the proportions of naive, central memory (TCM), effector memory (TEM), and terminally differentiated effector memory RA (TEMRA) T cells.

### Statistics

All statistical calculations were performed using GraphPad Prism (Versions 8.3-10). Binding affinities in the SPR drug competition assays were determined using three-parameter nonlinear regression with the equation *Y = Bottom + (Top - Bottom) / (1 + (X / IC50))*. Significance for killing assays or *in vivo* experiments was calculated by 2-way ANOVA for multiple comparisons. Exact p-values are shown in the figures. The representative data from cell assays are presented as individual values overlayed with bars of mean values and error bars for s.d. or s.e.m. as indicated in the legend. For primary T-cells, “n” refers to the number of human donors. For Jurkat and HEK293T cells, “n = 3” represents biological replicates, which are defined as different wells on the same plate. IC_50_ values in cell assays were calculated using three-parameter nonlinear regression. T_1/2_ values were determined using the equation *Y = (Y₀ - Plateau) × exp(-K × X) + Plateau* for one-phase decay analysis. All source data is attached as a source data file.

## Acknowledgements

This work was generously supported by Ludwig Cancer Research, the Swiss National Science Foundation (SNSF# 310030_204326), and the Carigest, Prostate Cancer, and ISREC Foundations to MI. The SNSF, the NCCR in Chemical Biology and in Molecular Systems Engineering, an ERC starting grant (#716058), and the Swiss Cancer League (KFS-5032-02-2020) supported BC. The Strategic Focus Area “Personalized Health and Related Technologies (#2021-446)” of the ETH Domain and the Anniversary Foundation of Swiss Life for Public Health and Medical Research supported LS. The NCCR Molecular Systems Engineering supported STR. We sincerely thank Prof. George Coukos for his support of our work. The anti-EpCAM scFv C215 sequence was kindly provided by Dr. Ludger Gross-Hovest. We wish to thank members of the Flow Cytometry Platform and the Animal Care Facility of the Epalinges Campus of the UNIL for their excellent support of our research.

## Data Availability

All data is available in the main text or the supplementary information materials. Source Data are provided with this paper. Plasmids are available on request.

## Author Contributions

M.I. and B.E.C. directed the study. L.S., G.G.A., S.S., and M.I. developed the project. L.S., G.G.A., R.C.R., R.R., M.T., E.C., A.M., and S.G. designed and performed the experiments. L.S., G.G.A., R.C.R., R.R., S.S., S.R., B.E.C., and M.I. analyzed the results. H.A. and M.B. collaborated on the avidity experiments. L.S. wrote the first draft of the manuscript with further writing and editing by M.I. All the authors discussed findings and provided feedback on the manuscript.

## References

1. Cappell, K.M. & Kochenderfer, J.N. Long-term outcomes following CAR T cell therapy: what we know so far. Nat Rev Clin Oncol 20, 359–371 (2023).

2. Siegel, R.L., Giaquinto, A.N. & Jemal, A. Cancer statistics, 2024. CA Cancer J Clin 74, 12–49 (2024).

3. Flugel, C.L. et al. Overcoming on-target, off-tumour toxicity of CAR T cell therapy for solid tumours. Nat Rev Clin Oncol 20, 49–62 (2023).

4. Lanitis, E., Coukos, G. & Irving, M. All systems go: converging synthetic biology and combinatorial treatment for CAR-T cell therapy. Curr Opin Biotechnol 65, 75–87 (2020).

5. Giordano Attianese, G.M.P., Ash, S. & Irving, M. Coengineering specificity, safety, and function into T cells for cancer immunotherapy. Immunol Rev (2023).

6. Weber, E.W. et al. Transient rest restores functionality in exhausted CAR-T cells through epigenetic remodeling. Science 372 (2021).

7. Wolf, B. et al. Safety and Tolerability of Adoptive Cell Therapy in Cancer. Drug Saf (2019).

8. Moghanloo, E. et al. Remote controlling of CAR-T cells and toxicity management: Molecular switches and next generation CARs. Translational Oncology 14, 101070 (2021).

9. Caruso, H.G. et al. Tuning Sensitivity of CAR to EGFR Density Limits Recognition of Normal Tissue While Maintaining Potent Antitumor Activity. Cancer Research 75, 3505–3518 (2015).

10. Park, S. et al. Micromolar affinity CAR T cells to ICAM-1 achieves rapid tumor elimination while avoiding systemic toxicity. Sci Rep 7, 14366 (2017).

11. Arcangeli, S. et al. Balance of Anti-CD123 Chimeric Antigen Receptor Binding Affinity and Density for the Targeting of Acute Myeloid Leukemia. Molecular Therapy 25, 1933–1945 (2017).

12. Di Roberto, R.B. et al. A Functional Screening Strategy for Engineering Chimeric Antigen Receptors with Reduced On-Target, Off-Tumor Activation. Molecular Therapy 28, 2564–2576 (2020).

13. Rafiq, S., Hackett, C.S. & Brentjens, R.J. Engineering strategies to overcome the current roadblocks in CAR T cell therapy. Nat Rev Clin Oncol 17, 147–167 (2020).

14. Wu, C.Y., Roybal, K.T., Puchner, E.M., Onuffer, J. & Lim, W.A. Remote control of therapeutic T cells through a small molecule-gated chimeric receptor. Science 350, aab4077 (2015).

15. Labanieh, L. et al. Enhanced safety and efficacy of protease-regulated CAR-T cell receptors. Cell 185, 1745–1763 e1722 (2022).

16. Li, H.S. et al. High-performance multiplex drug-gated CAR circuits. Cancer Cell 40, 1294–1305 e1294 (2022).

17. Jan, M. et al. Reversible ON- and OFF-switch chimeric antigen receptors controlled by lenalidomide. Sci. Transl. Med. 13, eabb6295 (2021).

18. Park, S. et al. Direct control of CAR T cells through small molecule-regulated antibodies. Nat Commun 12, 710 (2021).

19. Hill, Z.B., Martinko, A.J., Nguyen, D.P. & Wells, J.A. Human antibody-based chemically induced dimerizers for cell therapeutic applications. Nat Chem Biol 14, 112–117 (2018).

20. Kotla, V. et al. Mechanism of action of lenalidomide in hematological malignancies. J Hematol Oncol 2, 36 (2009).

21. Ludwig, H. et al. Prevention and management of adverse events of novel agents in multiple myeloma: a consensus of the European Myeloma Network. Leukemia 32, 1542–1560 (2018).

22. Giordano-Attianese, G. et al. A computationally designed chimeric antigen receptor provides a small-molecule safety switch for T-cell therapy. Nat Biotechnol (2020).

23. Giordano-Attianese, G. et al. A computationally designed chimeric antigen receptor provides a small-molecule safety switch for T-cell therapy. Nat Biotechnol (2020).

24. Cho, J.H., Collins, J.J. & Wong, W.W. Universal Chimeric Antigen Receptors for Multiplexed and Logical Control of T Cell Responses. Cell 173, 1426–1438 e1411 (2018).

25. Nixdorf, D. et al. Adapter CAR T cells to counteract T-cell exhaustion and enable flexible targeting in AML. Leukemia 37, 1298–1310 (2023).

26. Cartellieri, M. et al. Switching CAR T cells on and off: a novel modular platform for retargeting of T cells to AML blasts. Blood Cancer J 6, e458 (2016).

27. Zajc, C.U. et al. A conformation-specific ON-switch for controlling CAR T cells with an orally available drug. Proc Natl Acad Sci U S A 117, 14926–14935 (2020).

28. Shui, S. et al. A rational blueprint for the design of chemically-controlled protein switches. Nat Commun 12, 5754 (2021).

29. Garcia-Aranda, M., Perez-Ruiz, E. & Redondo, M. Bcl-2 Inhibition to Overcome Resistance to Chemo- and Immunotherapy. Int J Mol Sci 19 (2018).

30. Gong, M.C., Chang, S.S., Sadelain, M., Bander, N.H. & Heston, W.D. Prostate-specific membrane antigen (PSMA)-specific monoclonal antibodies in the treatment of prostate and other cancers. Cancer Metastasis Rev 18, 483–490 (1999).

31. Deeks, E.D. Venetoclax: First Global Approval. Drugs 76, 979–987 (2016).

32. Roberts, A.W. et al. Targeting BCL2 with Venetoclax in Relapsed Chronic Lymphocytic Leukemia. N Engl J Med 374, 311–322 (2016).

33. Tong, C. et al. Optimized tandem CD19/CD20 CAR-engineered T cells in refractory/relapsed B-cell lymphoma. Blood 136, 1632–1644 (2020).

34. Bachmann, M. The UniCAR system: A modular CAR T cell approach to improve the safety of CAR T cells. Immunol Lett 211, 13–22 (2019).

35. Cui, Y. et al. Construction and application of service quality evaluation system in the preclinical research on cardiovascular implant devices. BMC Med Inform Decis Mak 19, 37 (2019).

36. Souers, A.J. et al. ABT-199, a potent and selective BCL-2 inhibitor, achieves antitumor activity while sparing platelets. Nat Med 19, 202–208 (2013).

37. Bojar, D., Scheller, L., Hamri, G.C.-E., Xie, M. & Fussenegger, M. Caffeine-inducible gene switches controlling experimental diabetes. Nat Commun 9, 2318 (2018).

38. Scheller, L., Strittmatter, T., Fuchs, D., Bojar, D. & Fussenegger, M. Generalized extracellular molecule sensor platform for programming cellular behavior. Nat Chem Biol 14, 723–729 (2018).

39. Green, A.A. et al. Complex cellular logic computation using ribocomputing devices. Nature 548, 117–121 (2017).

40. Santoro, S.P. et al. T Cells Bearing a Chimeric Antigen Receptor against Prostate-Specific Membrane Antigen Mediate Vascular Disruption and Result in Tumor Regression. Cancer immunology research (2014).

41. Irving, M., Lanitis, E., Migliorini, D., Ivics, Z. & Guedan, S. Choosing the Right Tool for Genetic Engineering: Clinical Lessons from Chimeric Antigen Receptor-T Cells. Hum Gene Ther 32, 1044–1058 (2021).

42. Ayala Ceja, M., Khericha, M., Harris, C.M., Puig-Saus, C. & Chen, Y.Y. CAR-T cell manufacturing: Major process parameters and next-generation strategies. J Exp Med 221 (2024).

43. Ho, J.Y. et al. Promoter usage regulating the surface density of CAR molecules may modulate the kinetics of CAR-T cells in vivo. Mol Ther Methods Clin Dev 21, 237–246 (2021).

44. Grosse-Hovest, L. et al. Tumor-growth inhibition with bispecific antibody fragments in a syngeneic mouse melanoma model: the role of targeted T-cell co-stimulation via CD28. Int J Cancer 80, 138–144 (1999).

45. Birkinshaw, R.W. et al. Structures of BCL-2 in complex with venetoclax reveal the molecular basis of resistance mutations. Nat Commun 10, 2385 (2019).

46. Li, W. et al. Chimeric Antigen Receptor Designed to Prevent Ubiquitination and Downregulation Showed Durable Antitumor Efficacy. Immunity 53, 456–470 e456 (2020).

47. Zhu, I. et al. Modular design of synthetic receptors for programmed gene regulation in cell therapies. Cell 185, 1431–1443.e1416 (2022).

48. Cho, J.H. et al. Engineering advanced logic and distributed computing in human CAR immune cells. Nat Commun 12, 792 (2021).

49. Manhas, J., Edelstein, H.I., Leonard, J.N. & Morsut, L. The evolution of synthetic receptor systems. Nat Chem Biol (2022).

50. Tong, C. et al. Optimized tandem CD19/CD20 CAR-engineered T cells in refractory/relapsed B cell lymphoma. Blood, blood.2020005278 (2020).

51. Mirdita, M. et al. ColabFold: making protein folding accessible to all. Nat Methods 19, 679–682 (2022).

52. Marchand, A. et al. Rational Design of Chemically Controlled Antibodies and Protein Therapeutics. ACS Chem. Biol. (2023).

