## Supplemental Tables and Figures for "Drug-controlled CAR-T cells through the regulation of cell-cell interactions"

### Supplementary Material


Supplementary Table 1: Description of all plasmids used in the study ____ 2

Supplementary Table 2: Protein Sequences ____ 3

Supplementary Table 3: RNA and DNA sequences for genome editing ____ 7

Supplementary Table 4: Deep sequencing results after library sorts ____ 8

Supplementary Figure 1: Proof-of-principle for cell-surface expression of the DROP-CAR and dissociation of the R-domain from the S-chain by venetoclax. ____ 12

Supplementary Figure 2: In Silico Site Saturation Mutagenesis. ____ 13

Supplementary Figure 3: Library generation and DROP-CAR functionality sorts. ____ 14

Supplementary Figure 4: Drug sensitivity screening. ____ 15

Supplementary Figure 5: Characterization of enriched DROP-CAR (LD3) variants. ____ 16

Supplementary Figure 6: Alignment of varianats with Human Apolipoprotein E4 (ApoE4). 17

Supplementary Figure 7: Characterization of LD3 variants enriched from the library screening and evaluation of optimized DROP-CAR expression and function in Jurkat cells.18

Supplementary Figure 8: Cell avidity analysis with hinge cells counted as bound ____ 19

Supplementary Figure 9: Proof of concept for dual DROP-CARs ____ 20

Supplementary Figure 10: Adapting the DROP design to the generalized extracellular molecule sensor (GEMS) cytokine receptor platform. ____ 21

Supplementary Figure 11: DROP-CAR expression in primary human T Cells. ____ 22

Supplementary Figure 12: Tumor control by DROP-CAR T cells. ____ 24

### Supplementary Table 1: Description of all plasmids used in the study

| Name | Description | Figure |
|---|---|---|
| pVSV-G | VSV glycoprotein expression plasmid | Figure 3, Figure 4, S7, S8, S9, S11, S12 |
| R874 | Rev and Gag/Pol expression plasmid | Figure 3, Figure 4, S7, S8, S9, S11, S12 |
| pLS400_DROP_WT | Template plasmid for making the PCR with homology arms to integrate the initial DROP CAR into B3Z cells. | S1, Figure 2 |
| pAM193_pHLSec_LD3_A130G-R134V | Expression plasmid (CMV enhancer with a chicken B-actin promoter) for the mutated LD3 protein A130G; R134V | S7a |
| pAM191_pHLSec_LD3_A139R | Expression plasmid (CMV enhancer with a chicken B-actin promoter) for the mutated LD3 protein A139R | S7a |
| pLS147_secLD3-caff | Secretion of LD3-caffeine-nanobody fusion protein for assembly on the GEMS receptors (SV40 promoter) | S10 |
| pLS912_secLD3-A130G-R134V-caff | Secretion of LD3-A130G-R134V-caffeine-nanobody fusion protein for assembly on the GEMS receptors (SV40 promoter) | S10 |
| pLS392_STAT3 | STAT3 expression plasmid (SV40 promoter) | S10 |
| pLS858_Nanoluc | STAT3 dependent nanoluciferase reporter plasmid (STAT3 binding sites – minimal CMV promoter – Nanoluciferase-Fc) | S10 |
| S184-Bcl-xL-GEMS | Bcl-xL-GEMS receptor (SV40 promoter), developed by Shui et al. (2021) https://doi.org/10.1038/s41467-021-25735-9 | S10 |
| S193-Bcl2-GEMS | Bcl2-GEMS receptor (SV40 promoter), developed by Shui et al. (2021) https://doi.org/10.1038/s41467-021-25735-9 | S10 |
| pLSX135_dual-DROP-aCD19-Bcl-xL-aPSMA-Bcl-2 | Lentiviral vector (EF1α Promoter – ORF – WPRE) for expressing the optimized dual-DROP-CAR | S9 |
| pLSX052_DROP-CAR_A130G-R134V | Lentiviral vector (EF1α Promoter – ORF – WPRE) for expressing the optimized PSMA-DROP-CAR | Figure 3, Figure 4, S11, S12 |
| pGGA001_pELNS-EpCAM-DROP-CAR | Lentiviral vector (EF1α Promoter – ORF – WPRE) for expressing the optimized EpCAM-DROP-CAR | Figure 4, S11 |

### Supplementary Table 2: Protein Sequences

| Name | Description | Sequence |
|---|---|---|
| LD3 WT | The original LD3 protein developed in Giordano-Attianese & Gainza et al. 2020, https://doi.org/10.1038/s41587-019-0403-9 | GQRWELALGRFLEYLSWVSTLSEQVQEEL LSSQVTQELRALMDETMKELKAYKSELEEQ LTPVAEETRARLSKELQAAQARLGADMED VRGRLVQYRGEVQAMLGQSTEELRVRLAS HLIALQLRLIGDAFDLQKRLAVYQAGA |
| LD3 A130G; R134V | The most enriched variant during sorting and the basis for the final DROP-CAR | GQRWELALGRFLEYLSWVSTLSEQVQEEL LSSQVTQELRALMDETMKELKAYKSELEEQ LTPVAEETRARLSKELQAAQARLGADMED VRGRLVQYRGEVQAMLGQSTEELRVRLAS HLIGLQLVLIGDAFDLQKRLAVYQAGA |
| LD3 A139R | Another partially enriched variant, not chosen for the DROP-CAR | GQRWELALGRFLEYLSWVSTLSEQVQEEL LSSQVTQELRALMDETMKELKAYKSELEEQ LTPVAEETRARLSKELQAAQARLGADMED VRGRLVQYRGEVQAMLGQSTEELRVRLAS HLIALQLRLIGDRFDLQKRLAVYQAGA |
| LD3 Q132L | Another partially enriched variant, not chosen for the DROP-CAR | GQRWELALGRFLEYLSWVSTLSEQVQEEL LSSQVTQELRALMDETMKELKAYKSELEEQ LTPVAEETRARLSKELQAAQARLGADMED VRGRLVQYRGEVQAMLGQSTEELRVRLAS HLIALLLRLIGDAFDLQKRLAVYQAGA |
| Bcl-2 | Bcl-2 sequence based on PDB 4LVT. Used in the plasmids pLS400 and S193. | AHPGRTGYDNREIVMKYIHYKLSQRGYEW DAGDDVEENRTEAPEGTESEVVHLTLRQA GDDFSRRYRRDFAEMSSQLHLTPFTARGR FATVVEELFRDGVNWGRIVAFFEFGGVMC VESVNREMSPLVDNIALWMTEYLNRHLHT WIQDNGGWDAFVELYGP |
| Bcl-$2_{opt}$ | Bcl-2 sequence optimized for better solubility by Lajoie et al. 2020 https://doi.org/10.1126/science.aba6527 Used in the plasmids pLSX052, pLSX135, and pGGA001_pELNS-EpCAM-DROP-CAR. | AHAGRTGYDNREIVMKYIHYKLSQRGYEW DAGDDAEENRTEAPEGTESEVVHRALRDA GDDFERRYRRDFAEMSSQLHLTPDTARQR FETVVEELFRDGVNWGRIVAFFEFGGVMC VESVNREMSPLVDNIAEWMTEYLNRHLHT WIQDNGGWDAFVELYGP |
| Bcl-xL | Bcl-xL sequence used for dual DROP-CARs (pLSX135_dual-DROP-aCD19-Bcl-xL-aPSMA-Bcl-2) and DROP-GEMS (S184-Bcl-xL-GEMS). | MSQSNRELVVDFLSYKLSQKGYSWSQFSD VEENRTEAPEGTESEAVKQALREAGDEFEL RYRRAFSDLTSQLHITPGTAYQSFEQVVNE LFRDGVNWGRIVAFFSFGGALCVESVDKE MQVLVSRIAAWMATYLNDHLEPWIQENGG WDTFVELYGNNAAAESRKGQER |
| PZ1 scFv | scFv for the PZ1 DROP-CAR. | EVQLQQSGPELVKPGTSVRISCKTSGYTFT EYTIHWVKQSHGKSLEWIGNINPNNGGTTY NQKFEDKATLTVDKSSSTAYMELRSLTSED SAVYYCAAGWNFDYWGQGTTVTVSSGGG GSGGGGSGGGGSDIVMTQSHKFMSTSVG DRVSIICKASQDVGTAVDWYQQKPGQSPK LLIYWASTRHTGVPDRFTGSGSGTDFTLTIT NVQSEDLADYFCQQYNSYPLTFGAGTMLD LKR |
| Her2 scFv | scFv for the WT DROP-CAR for screening in BZ1 cells. | DIQMTQSPSSLSASVGDRVTITCRASQDVN TAVAWYQQKPGKAPKLLIYSASFLYSGVPS RFSGSRSGTDFTLTISSLQPEDFATYYCQQ HYTTPPTFGQGTKVEIKRTGSTSGSGKPGS |

| | | |
|---|---|---|
| | | GEGSEVQLVESGGGLVQPGGSLRLSCAASGFNIKDTYIHWVRQAPGKGLEWVARIYPTNGYTRYADSVKGRFTISADTSKNTAYLQMNSLRAEDTAVYYCSRWGGDGFYAMDVWGQGTLVTVSS |
| CD19 scFv | scFv used for dual DROP-CARs (pLSX135_dual-DROP-aCD19-Bcl-xL-aPSMA-Bcl-2). | DIQMTQTTSSLSASLGDRVTISCRASQDISKYLNWYQQKPDGTVKLLIYHTSRLHSGVPSRFSGSGSGTDYSLTISNLEQEDIATYFCQQGNTLPYTFGGGTKLEITGSTSGSGKPGSGEGSTKGEVKLQESGPGLVAPSQSLSVTCTVSGVSLPDYGVSWIRQPPRKGLEWLGVIWGSETTYYNSALKSRLTIIKDNSKSQVFLKMNSLQTDDTAIYYCAKHYYYGGSYAMDYWGQGTSVTVS |
| EpCAM scFv | scFv for the EpCAM DROP-CAR. | QVKLQQSGAELVRPGASVKLSCKASGYTFTNYWINWVKQRPGQGLEWIGNIYPSYIYTNYNQEFKDKVTLTVDESSSTAYMQLSSPTSEDSAVYYCTRSPYGYDEYGLDYWGQGTTVTVSSGGGGSGGGGSGGGGSDIELTQSPSSLTVTAGEKVTMNCKSSQSLLNSRNQKNYLTWYQQKPGQPPKLLIYWASTRESGVPDRFTGSGSGTDFTLTISSVQAEDLAVYYCQNDYVYPLTFGAGTKLEIKR |
| Bcl-2 GEMS | Bcl2-GEMS receptor (SV40 promoter, Plasmid: S193-Bcl2-GEMS), developed by Shui et al. (2021), used to test AND-NOT logic with co-secreted LD3-caffeine nanoboy fusions. Kappa chain secretion peptide, Bcl-2, EpoR ECD and TMD, IL6-ST ICD. Linkers uncolored. | MSETDTLLLWVLLLWVPGSTGDMAHPGRTGYDNREIVMKYIHYKLSQRGYEWDAGDDVEENRTEAPEGTESEVVHLTLRQAGDDFSRRYRRDFAEMSSQLHLTPFTARGRFATVVEELFRDGVNWGRIVAFFEFGGVMCVESVNREMSPLVDNIALWMTEYLNRHLHTWIQDNGGWDAFVELYGPTSGAPSPSLPDPKFESKAALLASRGSEELLCFTQRLEDLVCFWEEAASSGMDFNYSFSYQLEGESRKSCSLHQAPTVRGSVRFWCSLPTADTSSAVPLELQVTEASGSPRYHRIIHINEVVLLDAPAGLLARRAEEGSHVVLRWLPPPGAPMTTHIRYEVDVSAGNRAGGTQRVEVLEGRTECVLSNLRGGTRYTFAVRARMAEPSFSGFWSAWSEPASLLTASDLDPLILTLSLILVLISLLLTVLALLSAAANKRDLIKKHIWPNVPDPSKSHIAQWSPHTPPRHNFNSKDQMYSDGNFTDVSVVEIEANDKKPFPEDLKSLDLFKKEKINTEGHSSGIGGSSCMSSSRPSISSSDENESSQNTSSTVQASTVVHSGYRHQVPSVQVFSRSESTQPLLDSEERPEDLQLVDHVDGGDGILPRQQYFKQNCSQHESSPDISHFERSKQVSSVNEEDFVRLKQQISDHISQSCGSGQMKMFQEVSAADAFGPGTEGQVERFETVGMEAATDEGMPKSYLPQTVRQGGYMPQASTGV* |
| LD3-anti-caff fusion protein | LD3-anti-caff fusion protein (SV40 promoter, Plasmid: pLS147_secLD3-WT-caff), used to test AND-NOT logic with Bcl-2-GEMS. Albumin secretion peptide, LD3, anti-caffeine VHH (PDB 6QTL). Linkers uncolored. | MKWVTFISLLFLFSSAYSTGTSGQRWELALGRFLEYLSWVSTLSEQVQEELLSSQVTQELRALMDETMKELKAYKSELEEQLTPVAEETRARLSKELQAAQARLGADMEDVRGRLVQYRGEVQAMLGQSTEELRVRLASHLIALQLRLIGDAFDLQKRLAVYQAGAGASGSQVQLVESGGGLVQAGGSLRLSCTASGRTGTIYSMAWFRQAPGKEREFLATVGWSSGITYYMDSVKG |

| | | |
|---|---|---|
| | | RFTISRDKGKNTVYLQMDSLKPEDTAVYYC<br>TATRAYSVGYDYWGQGTQVTVSSR* |
| DROP_WT | WT-DROP CAR sequence before mutations. Expressed genomically from the murine BZ1 T-cell line. (template plasmid: pLS400_DROP_WT).<br>Kappa chain secretion peptide, LD3-WT, Strep tag, mCD28 hinge, mCD28 TMD, mCD28 ICD (LL:GG mutant), mCD3Z, P2A, Her2 scFv (4D5), Bcl-2, V5. Linkers uncolored. | METDTLLLWVLLLWVPGSTGDTGQRWELA<br>LGRFLEYLSWVSTLSEQVQEELLSSQVTQE<br>LRALMDETMKELKAYKSELEEQLTPVAEET<br>RARLSKELQAAQARLGADMEDVRGRLVQY<br>RGEVQAMLGQSTEELRVRLASHLIALQLRLI<br>GDAFDLQKRLAVYQAGAAESANWSHPQFE<br>KGGGGSGGGGSNWSHPQFEKSAIEFMYP<br>PPYLDNERSNGTIIHIKEKHLCHTQSSPKLF<br>WALVVVAGVLFCYGLLVTVALCVIWTNSRR<br>NRGGQSDYMNMTPRRPGLTRKPYQPYAP<br>ARDFAAYRPRAKFSRSAETAANLQDPNQL<br>YNELNLGRREEYDVLEKKRARDPEMGGKQ<br>QRRRNPQEGVYNALQKDKMAEAYSEIGTK<br>GERRRGKGHDGLYQGLSTATKDTYDALHM<br>QTLAPRGGGGSGATNFSLLKQAGDVEENP<br>GPGGGMETDTLLLWVLLLWVPGSTGDASD<br>IQMTQSPSSLSASVGDRVTITCRASQDVNT<br>AVAWYQQKPGKAPKLLIYSASFLYSGVPSR<br>FSGSRSGTDFTLTISSLQPEDFATYYCQQH<br>YTTPPTFGQGTKVEIKRTGSTSGSGKPGSG<br>EGSEVQLVESGGGLVQPGGSLRLSCAASG<br>FNIKDTYIHWVRQAPGKGLEWVARIYPTNG<br>YTRYADSVKGRFTISADTSKNTAYLQMNSL<br>RAEDTAVYYCSRWGGDGFYAMDVWGQG<br>TLVTVSSGGPGMAHPGRTGYDNREIVMKYI<br>HYKLSQRGYEWDAGDDVEENRTEAPEGT<br>ESEVVHLTLRQAGDDFSRRYRRDFAEMSS<br>QLHLTPFTARGRFATVVEELFRDGVNWGRI<br>VAFFEFGGVMCVESVNREMSPLVDNIALW<br>MTEYLNRHLHTWIQDNGGWDAFVELYGPG<br>GGSGKPIPNPLLGLDST* |
| PZ1-DROP-CAR | DROP-CAR expressed from the lentiviral pELNS vector (pLSX052_DROP-CAR_A130G-R134V).<br>CD8 secretion signal, FLAG, LD3 A130G; R134V, hCD8 hinge, hCD28 TMD, hCD28 ICD, hCD3Z, T2A, CD8 secretion signal, PZ1 scFv, Bcl2$_{opt}$, myc. Linkers uncolored. | MALPVTALLLPLALLLHAARPGSDYKDDDD<br>KGQRWELALGRFLEYLSWVSTLSEQVQEE<br>LLSSQVTQELRALMDETMKELKAYKSELEE<br>QLTPVAEETRARLSKELQAAQARLGADME<br>DVRGRLVQYRGEVQAMLGQSTEELRVRLA<br>SHLIGLQLVLIGDAFDLQKRLAVYQAGAAST<br>TTPAPRPPTPAPTIASQPLSLRPEACRPAA<br>GGAVHTRGLDFACDFWVLVVVGGVLACYS<br>LLVTVAFIIFWVRSKRSRLLHSDYMNMTPR<br>RPGPTRKHYQPYAPPRDFAAYRSRVKFSR<br>SADAPAYQQGQNQLYNELNLGRREEYDVL<br>DKRRGRDPEMGGKPRRKNPQEGLYNELQ<br>KDKMAEAYSEIGMKGERRRGKGHDGLYQ<br>GLSTATKDTYDALHMQALPPRSGGSGEGR<br>GSLLTCGDVEENPGPMALPVTALLLPLALLL<br>HAARPEVQLQQSGPELVKPGTSVRISCKTS<br>GYTFTEYTIHWVKQSHGKSLEWIGNINPNN<br>GGTTYNQKFEDKATLTVDKSSSTAYMELRS<br>LTSEDSAVYYCAAGWNFDYWGQGTTVTVS<br>SGGGGSGGGGSGGGGSDIVMTQSHKFMS<br>TSVGDRVSIICKASQDVGTAVDWYQQKPG<br>QSPKLLIYWASTRHTGVPDRFTGSGSGTDF<br>TLTITNVQSEDLADYFCQQYNSYPLTFGAG<br>TMLDLKRGGPGAHAGRTGYDNREIVMKYI |

| | | |
|---|---|---|
| | | HYKLSQRGYEWDAGDDAEENRTEAPEGT<br>ESEVVHRALRDAGDDFERRYRRDFAEMSS<br>QLHLTPDTARQRFETVVEELFRDGVNWGRI<br>VAFFEFGGVMCVESVNREMSPLVDNIAEW<br>MTEYLNRHLHTWIQDNGGWDAFVELYGPS<br>MREQKLISEEDL* |
| CD19, PZ1 Dual-DROP-CAR | Dual DROP-CAR expressed from the lentiviral pELNS vector (pLSX135_dual-DROP-aCD19-Bcl-xL-aPSMA-Bcl-2). CD8 secretion signal, FLAG, LD3 A130G; R134V, hCD8 hinge, hCD28 TMD, hCD28 ICD, hCD3Z, T2A, CD8 secretion signal, V5, CD19 scFv, Bcl-xL, T2A, CD8 secretion signal, HA, PZ1 scFv, $Bcl2_{opt}$, myc. Linkers uncolored. | MALPVTALLLPLALLLHAARPGSDYKDDDD<br>KGQRWELALGRFLEYLSWVSTLSEQVQEE<br>LLSSQVTQELRALMDETMKELKAYKSELEE<br>QLTPVAEETRARLSKELQAAQARLGADME<br>DVRGRLVQYRGEVQAMLGQSTEELRVRLA<br>SHLIGLQLVLIGDAFDLQKRLAVYQAGAAST<br>TTPAPRPPTPAPTIASQPLSLRPEACRPAA<br>GGAVHTRGLDFACDFWVLVVVGGVLACYS<br>LLVTVAFIIFWVRSKRSRLLHSDYMNMTPR<br>RPGPTRKHYQPYAPPRDFAAYRSRVKFSR<br>SADAPAYQQGQNQLYNELNLGRREEYDVL<br>DKRRGRDPEMGGKPRRKNPQEGLYNELQ<br>KDKMAEAYSEIGMKGERRRGKGHDGLYQ<br>GLSTATKDTYDALHMQALPPRSGGSGEGR<br>GSLLTCGDVEENPGPMALPVTALLLPLALLL<br>HAARPGSKPIPNPLLGLDSTGGGDIQMTQT<br>TSSLSASLGDRVTISCRASQDISKYLNWYQ<br>QKPDGTVKLLIYHTSRLHSGVPSRFSGSGS<br>GTDYSLTISNLEQEDIATYFCQQGNTLPYTF<br>GGGTKLEITGSTSGSGKPGSGEGSTKGEV<br>KLQESGPGLVAPSQSLSVTCTVSGVSLPDY<br>GVSWIRQPPRKGLEWLGVIWGSETTYYNS<br>ALKSRLTIIKDNSKSQVFLKMNSLQTDDTAI<br>YYCAKHYYYGGSYAMDYWGQGTSVTVSG<br>GGGMSQSNRELVVDFLSYKLSQKGYSWS<br>QFSDVEENRTEAPEGTESEAVKQALREAG<br>DEFELRYRRAFSDLTSQLHITPGTAYQSFE<br>QVVNELFRDGVNWGRIVAFFSFGGALCVE<br>SVDKEMQVLVSRIAAWMATYLNDHLEPWI<br>QENGGWDTFVELYGNNAAAESRKGQERG<br>RKRRSGSGEGRGSLLTCGDVEENPGPMAL<br>PVTALLLPLALLLHAARPYPYDVPDYAGGG<br>SEVQLQQSGPELVKPGTSVRISCKTSGYTF<br>TEYTIHWVKQSHGKSLEWIGNINPNNGGTT<br>YNQKFEDKATLTVDKSSSTAYMELRSLTSE<br>DSAVYYCAAGWNFDYWGQGTTVTVSSGG<br>GGSGGGGSGGGGSDIVMTQSHKFMSTSV<br>GDRVSIICKASQDVGTAVDWYQQKPGQSP<br>KLLIYWASTRHTGVPDRFTGSGSGTDFTLTI<br>TNVQSEDLADYFCQQYNSYPLTFGAGTML<br>DLKRGGPGAHAGRTGYDNREIVMKYIHYKL<br>SQRGYEWDAGDDAEENRTEAPEGTESEV<br>VHRALRDAGDDFERRYRRDFAEMSSQLHL<br>TPDTARQRFETVVEELFRDGVNWGRIVAFF<br>EFGGVMCVESVNREMSPLVDNIAEWMTEY<br>LNRHLHTWIQDNGGWDAFVELYGPSMRE<br>QKLISEEDL* |

### Supplementary Table 3: RNA and DNA sequences for genome editing

| **Type** | **Experiment description** | **Sequence** |
|---|---|---|
| crRNA – TCR beta locus | Targeting the TCR beta locus to integrate the initial DROP-CAR | ACCGAAGUACUGUUCAUAAU |
| crRNA – LD3 locus -1 | Targeting the LD3 site to introduce a frameshift mutation | GCCCUACAACUGCGUCUGAU |
| crRNA – LD3 clone F6-1 | Targeting the LD3 site to repair the frameshift mutation | CACCGAUCAGACGCAGUUGU |
| crRNA – LD3 clone F6-2 | Targeting the LD3 site to repair the frameshift mutation | ACCGAUCAGACGCAGUUUGU |
| crRNA – LD3 clone C8-1 | Targeting the LD3 site to repair the frameshift mutation | CCCUACAAACUGCGUCUGAU |
| crRNA – LD3 clone C8-2 | Targeting the LD3 site to repair the frameshift mutation | UCUGAUUGCCCUACACUGAU |
| ssODN – NNK1 | Site saturation mutagenesis of A130 and R134 | CATGCTGGGCCAATCGACGGAAGAACTGCGTGTCCGCCTGGCGAGCCATCTGATT**NNK**CTACAACTG**NNK**CTGATCGGTGACGCATTCGACCTACAGAAACGCCTGGCTGTCTACCAAGCAGGCG |
| ssODN – NNK2 | Site saturation mutagenesis of Q132 | CATGCTGGGCCAATCGACGGAAGAACTGCGTGTCCGCCTGGCGAGCCATCTGATTGCCCTA**NNK**CTGCGTCTGATCGGTGACGCATTCGACCTACAGAAACGCCTGGCTGTCTACCAAGCAGGCG |
| ssODN – NNK3 | Site saturation mutagenesis of D138 | GGAAGAACTGCGTGTCCGCCTGGCGAGCCATCTGATTGCCCTACAACTGCGTCTGATCGGT**NNK**GCATTCGACCTACAGAAACGCCTGGCTGTCTACCAAGCAGGCGCTGCTGAGAGCGCTAATT |
| ssODN – NNK4 | Site saturation mutagenesis of A139 | AGAACTGCGTGTCCGCCTGGCGAGCCATCTGATTGCCCTACAACTGCGTCTGATCGGTGAC**NNK**TTCGACCTACAGAAACGCCTGGCTGTCTACCAAGCAGGCGCTGCTGAGAGCGCTAATTGGA |
| LD3_oligo_F | PCR primer for genomic LD3 amplification (forward) | AAAGAACTACAGGCCGCACA |
| LD3_oligo_R | PCR primer for genomic LD3 amplification (reverse) | CGAACTGAGGGTGTGACCAG |

**Supplementary Table 4: Deep sequencing results after library sorts**

Enriched variants are marked in red, undetected variants are marked in grey. Wild type residues are excluded from analysis. For the double mutant library NNK1, left to right refers to mutations of A130 and top to bottom to mutations of R134.

| V5/Strep double positive (assembled CARs; sort 1) | | | | | | | | | | | | | | | | | | | | | |
|---|---|---|---|---|---|---|---|---|---|---|---|---|---|---|---|---|---|---|---|---|---|
| | | A | R | N | D | C | Q | E | G | H | I | L | K | M | F | P | S | T | W | Y | V |
| **NNK1 A130X R134X** | A | 0.1352 | 0.0165 | 0 | 0 | 0 | 0 | 0.0002 | 0.4474 | 0 | 0 | 0 | 0 | 0 | 0.0019 | 0.0004 | 0.0068 | 0 | 0.0023 | 0.0015 | 0.0006 |
| | R | X | 0.0076 | 0 | 0.0038 | 0.0118 | 0.0004 | 0.0343 | 1.3922 | 0 | 0.0059 | 0.0009 | 0 | 0.0002 | 0.0006 | 0.1502 | 0.3079 | 0.0133 | 0.0205 | 0.0064 | 0.0631 |
| | N | 0 | 0 | 0 | 0 | 0 | 0 | 0 | 0.0004 | 0 | 0 | 0 | 0 | 0 | 0 | 0 | 0.0002 | 0 | 0 | 0 | 0 |
| | D | 0 | 0 | 0 | 0 | 0.0055 | 0 | 0 | 0 | 0 | 0 | 0 | 0 | 0 | 0 | 0 | 0.0019 | 0 | 0 | 0 | 0 |
| | C | 0.1678 | 0.0017 | 0.0040 | 0 | 0.0002 | 0 | 0.0004 | 0.3646 | 0 | 0 | 0 | 0 | 0 | 0 | 0.0055 | 0.0009 | 0 | 0.0002 | 0.0040 | 0.0038 |
| | Q | 0.0148 | 0 | 0 | 0.0002 | 0 | 0 | 0 | 0.0349 | 0 | 0 | 0 | 0 | 0 | 0 | 0.0099 | 0.0002 | 0 | 0 | 0 | 0 |
| | E | 0.0002 | 0 | 0 | 0 | 0.0004 | 0 | 0 | 0.0006 | 0 | 0 | 0 | 0 | 0 | 0 | 0 | 0 | 0 | 0 | 0 | 0 |
| | G | 0.2491 | 0.0055 | 0 | 0.0083 | 0.0076 | 0 | 0.0002 | 0.7957 | 0.0002 | 0.0085 | 0.0027 | 0 | 0.0021 | 0 | 0.0178 | 0.0011 | 0.0002 | 0 | 0 | 0.0008 |
| | H | 0.0142 | 0.0002 | 0 | 0 | 0 | 0.0006 | 0 | 0.0851 | 0 | 0 | 0.0082 | 0 | 0 | 0 | 0.0002 | 0.0053 | 0 | 0.0104 | 0 | 0 |
| | I | 0.0463 | 0 | 0 | 0.0004 | 0.0104 | 0 | 0 | 0.0008 | 0 | 0 | 0 | 0 | 0 | 0 | 0 | 0 | 0 | 0 | 0.0013 | 0.0002 |
| | L | 0.0169 | 0 | 0 | 0.0013 | 0.0019 | 0.0002 | 0.0068 | 0.0131 | 0 | 0.0021 | 0 | 0 | 0 | 0.0085 | 0 | 0.0193 | 0 | 0 | 0.0002 | 0.0080 |
| | K | 0.1253 | 0 | 0 | 0 | 0.0011 | 0 | 0.0002 | 0.2495 | 0 | 0 | 0.0038 | 0 | 0 | 0.0002 | 0.0436 | 0.0728 | 0 | 0 | 0 | 0.0222 |
| | M | 0.0468 | 0 | 0.0030 | 0 | 0 | 0.0074 | 0.0002 | 0.0844 | 0 | 0 | 0.0013 | 0 | 0 | 0 | 0.0015 | 0.0017 | 0 | 0 | 0 | 0 |
| | F | 0.0102 | 0.0072 | 0 | 0 | 0 | 0 | 0 | 0 | 0 | 0.0108 | 0.0021 | 0 | 0 | 0 | 0 | 0 | 0 | 0 | 0 | 0.0104 |
| | P | 0.0009 | 0 | 0 | 0 | 0 | 0 | 0 | 0.0025 | 0 | 0 | 0 | 0 | 0.0017 | 0 | 0 | 0.0002 | 0 | 0 | 0 | 0 |
| | S | 0.0607 | 0.0017 | 0 | 0.0032 | 0 | 0 | 0.0008 | 1.0589 | 0 | 0.0044 | 0.0061 | 0 | 0.0038 | 0 | 0.0013 | 0.0004 | 0.0015 | 0.0180 | 0 | 0.0157 |
| | T | 0.0087 | 0.0195 | 0 | 0.0002 | 0 | 0 | 0.0066 | 0.1115 | 0 | 0 | 0 | 0 | 0.0017 | 0.0028 | 0.0004 | 0.0076 | 0 | 0 | 0 | 0 |
| | W | 0.0008 | 0.0159 | 0 | 0 | 0 | 0 | 0 | 0.0002 | 0 | 0 | 0 | 0 | 0.0082 | 0 | 0 | 0 | 0 | 0.0051 | 0 | 0 |
| | Y | 0.0002 | 0.0245 | 0 | 0 | 0 | 0 | 0 | 0 | 0 | 0 | 0 | 0 | 0 | 0 | 0 | 0 | 0 | 0 | 0 | 0 |
| | V | 0.0436 | 0 | 0 | 0 | 0 | 0.0114 | 0.0066 | 0.1086 | 0.0051 | 0 | 0 | 0 | 0 | 0 | 0.0006 | 0.0137 | 0 | 0 | 0.0004 | 0.0002 |
| **NNK2 Q132X** | | 3.5404 | 7.1066 | 1.4477 | 1.8442 | 1.8394 | X | 2.4766 | 4.8437 | 1.8318 | 1.3615 | 5.4871 | 2.2791 | 2.2658 | 2.0967 | 2.1395 | 5.1317 | 2.7119 | 1.7581 | 1.4409 | 4.1860 |
| **NNK3 D138X** | | 0.0525 | 0.0315 | 0.0315 | X | 0.0415 | 0.0193 | 0.2527 | 0.1232 | 0.0078 | 0.0034 | 0.0214 | 0 | 0.0133 | 0.0053 | 0 | 0.0610 | 0.0087 | 0.0140 | 0.0072 | 0.0387 |
| **NNK4 A139X** | | X | 0.6749 | 0.2389 | 0.4148 | 0.3832 | 0.2508 | 0.4448 | 1.2090 | 0.1594 | 0.3445 | 1.5209 | 0.1194 | 0.6992 | 0.5151 | 0.4323 | 1.4198 | 0.7811 | 0.7802 | 0.3949 | 1.0883 |

### Second selection of GFP positive cells (functional CARs; sort 3)

| | | A | R | N | D | C | Q | E | G | H | I | L | K | M | F | P | S | T | W | Y | V |
|---|---|---|---|---|---|---|---|---|---|---|---|---|---|---|---|---|---|---|---|---|---|
| **NNK1 A130X R134X** | A | 0.0017 | 0.0007 | 0 | 0 | 0 | 0 | 0 | 0.0705 | 0 | 0 | 0 | 0 | 0 | 0 | 0 | 0 | 0 | 0 | 0 | 0 |
| | R | X | 0 | 0 | 0.0056 | 0 | 0 | 0.0013 | 0.8391 | 0 | 0 | 0 | 0 | 0 | 0 | 0.0020 | 0.0053 | 0.0083 | 0.0003 | 0 | 0.0218 |
| | N | 0 | 0 | 0 | 0 | 0 | 0 | 0 | 0 | 0 | 0 | 0 | 0 | 0 | 0 | 0 | 0 | 0 | 0 | 0 | 0 |
| | D | 0 | 0 | 0 | 0 | 0 | 0 | 0 | 0 | 0 | 0 | 0 | 0 | 0 | 0 | 0 | 0 | 0 | 0 | 0 | 0 |
| | C | 0.0242 | 0 | 0.0003 | 0 | 0 | 0 | 0 | 0.0119 | 0 | 0 | 0 | 0 | 0 | 0 | 0.0003 | 0 | 0 | 0 | 0 | 0 |
| | Q | 0.0003 | 0 | 0 | 0 | 0 | 0 | 0 | 0.0010 | 0 | 0 | 0 | 0 | 0 | 0 | 0 | 0 | 0 | 0 | 0 | 0 |
| | E | 0 | 0 | 0 | 0 | 0 | 0 | 0 | 0.0003 | 0 | 0 | 0 | 0 | 0 | 0 | 0 | 0 | 0 | 0 | 0 | 0 |
| | G | 0.0139 | 0 | 0 | 0 | 0.0003 | 0 | 0 | 0.0486 | 0 | 0 | 0 | 0 | 0 | 0 | 0 | 0 | 0 | 0 | 0 | 0 |
| | H | 0.0103 | 0 | 0 | 0 | 0 | 0.0007 | 0 | 0.0010 | 0 | 0 | 0.0003 | 0 | 0 | 0 | 0 | 0 | 0 | 0 | 0 | 0 |
| | I | 0 | 0 | 0 | 0 | 0 | 0 | 0 | 0 | 0 | 0 | 0 | 0 | 0 | 0 | 0 | 0 | 0 | 0 | 0 | 0 |
| | L | 0.0023 | 0 | 0 | 0 | 0 | 0 | 0 | 0.0007 | 0 | 0 | 0 | 0 | 0 | 0 | 0 | 0.0003 | 0 | 0 | 0 | 0 |
| | K | 0.0023 | 0 | 0 | 0 | 0 | 0 | 0 | 0.0298 | 0 | 0 | 0 | 0 | 0 | 0 | 0.0007 | 0.0040 | 0 | 0 | 0 | 0 |
| | M | 0.0003 | 0 | 0 | 0 | 0 | 0 | 0 | 0.0010 | 0 | 0 | 0 | 0 | 0 | 0 | 0 | 0 | 0 | 0 | 0 | 0 |
| | F | 0 | 0.0003 | 0 | 0 | 0 | 0 | 0 | 0 | 0 | 0 | 0 | 0 | 0 | 0 | 0 | 0 | 0 | 0 | 0 | 0 |
| | P | 0.0010 | 0 | 0 | 0 | 0 | 0 | 0 | 0.0063 | 0 | 0 | 0 | 0 | 0 | 0 | 0 | 0 | 0 | 0 | 0 | 0 |
| | S | 0.0705 | 0 | 0 | 0.0003 | 0 | 0 | 0.0013 | 2.2738 | 0 | 0 | 0 | 0 | 0 | 0 | 0 | 0 | 0 | 0 | 0 | 0.0010 |
| | T | 0.0007 | 0 | 0 | 0 | 0 | 0 | 0 | 0.0026 | 0 | 0 | 0 | 0 | 0 | 0 | 0 | 0 | 0 | 0 | 0 | 0 |
| | W | 0 | 0 | 0 | 0 | 0 | 0 | 0 | 0 | 0 | 0 | 0 | 0 | 0.0003 | 0 | 0 | 0 | 0 | 0 | 0 | 0 |
| | Y | 0 | 0 | 0 | 0 | 0 | 0 | 0 | 0 | 0 | 0 | 0 | 0 | 0 | 0 | 0 | 0 | 0 | 0 | 0 | 0 |
| | V | 0.0013 | 0 | 0 | 0 | 0 | 0 | 0 | 0.0308 | 0 | 0 | 0 | 0 | 0 | 0 | 0 | 0 | 0 | 0 | 0 | 0 |
| **NNK2 Q132X** | | 1.3407 | 4.3928 | 1.9270 | 1.1707 | 0.5939 | X | 2.3079 | 3.9997 | 0.5873 | 0.7230 | 5.8417 | 1.2114 | 4.0076 | 5.1981 | 2.7794 | 4.4712 | 2.5918 | 3.6049 | 0.9635 | 2.2265 |
| **NNK3 D138X** | | 0.0020 | 0.0003 | 0.0116 | X | 0 | 0 | 0.0083 | 0.0685 | 0.0003 | 0 | 0.0007 | 0 | 0 | 0 | 0 | 0.0007 | 0 | 0.0003 | 0.0017 | 0.0056 |
| **NNK4 A139X** | | X | 0.2601 | 1.8298 | 1.2653 | 1.5816 | 0.0725 | 0.1158 | 3.8217 | 0.0030 | 0.4583 | 4.1837 | 0.0026 | 0.0513 | 0.2842 | 0.0490 | 0.4586 | 1.0267 | 0.3349 | 1.3616 | 0.7001 |

### Second drug-sensitivity selection; sort 5

| | | A | R | N | D | C | Q | E | G | H | I | L | K | M | F | P | S | T | W | Y | V |
|---|---|---|---|---|---|---|---|---|---|---|---|---|---|---|---|---|---|---|---|---|---|
| **NNK1 A130X R134X** | A | 0.0472 | 0 | 0 | 0 | 0 | 0 | 0 | 0.0079 | 0 | 0 | 0 | 0 | 0 | 0 | 0 | 0 | 0 | 0 | 0 | 0 |
| | R | X | 0 | 0 | 0.0018 | 0 | 0 | 0.0240 | 0.0797 | 0 | 0 | 0 | 0 | 0 | 0 | 0.0011 | 0.0053 | 0.0061 | 0 | 0 | 0.0087 |
| | N | 0 | 0 | 0 | 0 | 0 | 0 | 0 | 0.0011 | 0 | 0 | 0 | 0 | 0 | 0 | 0 | 0 | 0 | 0 | 0 | 0 |
| | D | 0 | 0 | 0 | 0 | 0 | 0 | 0 | 0 | 0 | 0 | 0 | 0 | 0 | 0 | 0 | 0 | 0 | 0 | 0 | 0 |
| | C | 0.0061 | 0.0008 | 0.0003 | 0 | 0 | 0 | 0 | 0.1275 | 0 | 0 | 0 | 0 | 0 | 0 | 0 | 0 | 0 | 0 | 0 | 0 |
| | Q | 0 | 0 | 0 | 0 | 0 | 0 | 0 | 0 | 0 | 0 | 0 | 0 | 0 | 0 | 0 | 0 | 0 | 0 | 0 | 0 |
| | E | 0 | 0 | 0 | 0 | 0 | 0 | 0 | 0 | 0 | 0 | 0 | 0 | 0 | 0 | 0 | 0.0005 | 0 | 0 | 0 | 0 |
| | G | 0.0058 | 0 | 0 | 0.0011 | 0 | 0 | 0 | 0.2827 | 0 | 0 | 0 | 0 | 0 | 0 | 0 | 0 | 0 | 0 | 0 | 0 |

| | | | | | | | | | | | | | | | | | | | | | |
|---|---|---|---|---|---|---|---|---|---|---|---|---|---|---|---|---|---|---|---|---|---|
| | H | 0.0024 | 0 | 0 | 0 | 0 | 0 | 0 | 0 | 0 | 0 | 0 | 0 | 0 | 0 | 0 | 0 | 0 | 0.0003 | 0 | 0 |
| | I | 0.0005 | 0 | 0 | 0 | 0 | 0 | 0 | 0 | 0 | 0 | 0 | 0 | 0 | 0 | 0 | 0 | 0 | 0 | 0 | 0 |
| | L | 0.0013 | 0 | 0 | 0 | 0 | 0 | 0 | 0.0011 | 0 | 0 | 0 | 0 | 0 | 0.0003 | 0 | 0.0003 | 0 | 0 | 0 | 0 |
| | K | 0.0018 | 0 | 0 | 0 | 0 | 0 | 0 | 0.0011 | 0 | 0 | 0 | 0 | 0 | 0 | 0.0003 | 0.0457 | 0 | 0 | 0 | 0 |
| | M | 0.0003 | 0 | 0 | 0 | 0 | 0 | 0 | 0.0013 | 0 | 0 | 0 | 0 | 0 | 0 | 0 | 0 | 0 | 0 | 0 | 0 |
| | F | 0 | 0 | 0 | 0 | 0 | 0 | 0 | 0 | 0 | 0 | 0 | 0 | 0 | 0 | 0 | 0 | 0 | 0 | 0 | 0 |
| | P | 0.0003 | 0 | 0 | 0 | 0 | 0 | 0 | 0.0140 | 0 | 0 | 0 | 0 | 0 | 0 | 0 | 0 | 0 | 0 | 0 | 0 |
| | S | 0.0628 | 0.0037 | 0 | 0 | 0 | 0 | 0.0011 | 5.3266 | 0 | 0 | 0 | 0 | 0 | 0 | 0 | 0.0003 | 0 | 0.0003 | 0 | 0.0005 |
| | T | 0.0003 | 0 | 0 | 0 | 0 | 0 | 0 | 0.0058 | 0 | 0 | 0 | 0 | 0 | 0 | 0 | 0 | 0 | 0 | 0 | 0 |
| | W | 0 | 0 | 0 | 0 | 0 | 0 | 0 | 0 | 0 | 0 | 0 | 0 | 0 | 0 | 0 | 0 | 0 | 0 | 0 | 0 |
| | Y | 0 | 0.0003 | 0 | 0 | 0 | 0 | 0 | 0.0008 | 0 | 0 | 0 | 0 | 0 | 0 | 0 | 0 | 0 | 0 | 0 | 0 |
| | V | 0.0185 | 0 | 0 | 0.0003 | 0 | 0.0005 | 0 | 1.0879 | 0 | 0 | 0 | 0 | 0 | 0 | 0 | 0.0003 | 0 | 0 | 0 | 0 |
| **NNK2 Q132X** | | 2.5742 | 0.2227 | 0.3162 | 1.1216 | 2.7975 | X | 2.0944 | 7.4670 | 0.0852 | 1.8300 | 12.0227 | 0.0433 | 8.4300 | 5.4839 | 11.8844 | 8.1374 | 1.9150 | 3.5006 | 0.4231 | 4.4008 |
| **NNK3 D138X** | | 0.0003 | 0 | 0.0026 | X | 0.0003 | 0.0003 | 0.0032 | 0.0248 | 0.0003 | 0 | 0.0005 | 0 | 0.0003 | 0 | 0 | 0 | 0.0003 | 0 | 0 | 0.0013 |
| **NNK4 A139X** | | X | 2.2525 | 0.0185 | 0.0367 | 0.1180 | 0.0029 | 0.0238 | 3.6386 | 0.0018 | 0.0137 | 0.0953 | 0.0008 | 0.0090 | 0.2280 | 0.0892 | 0.0179 | 0.0322 | 0.0206 | 1.2650 | 0.0705 |

**Third drug-sensitivity selection; sort 6 - population V5-medium**

| | | A | R | N | D | C | Q | E | G | H | I | L | K | M | F | P | S | T | W | Y | V |
|---|---|---|---|---|---|---|---|---|---|---|---|---|---|---|---|---|---|---|---|---|---|
| **NNK1 A130X R134X** | A | 0.0541 | 0 | 0 | 0 | 0 | 0 | 0.0004 | 0.0021 | 0 | 0 | 0 | 0 | 0 | 0 | 0 | 0 | 0 | 0 | 0 | 0 |
| | R | X | 0 | 0 | 0.0017 | 0 | 0 | 0.0107 | 0.0689 | 0 | 0 | 0 | 0 | 0 | 0 | 0 | 0.0021 | 0.0017 | 0 | 0 | 0.0017 |
| | N | 0 | 0 | 0 | 0 | 0 | 0 | 0 | 0.0004 | 0 | 0 | 0 | 0 | 0 | 0 | 0 | 0 | 0 | 0 | 0 | 0 |
| | D | 0 | 0 | 0 | 0 | 0 | 0 | 0 | 0 | 0 | 0 | 0 | 0 | 0 | 0 | 0 | 0 | 0 | 0 | 0 | 0 |
| | C | 0.0012 | 0 | 0 | 0 | 0 | 0 | 0 | 0 | 0 | 0 | 0 | 0 | 0 | 0 | 0 | 0 | 0 | 0 | 0 | 0 |
| | Q | 0 | 0 | 0 | 0 | 0 | 0 | 0 | 0 | 0 | 0 | 0 | 0 | 0 | 0 | 0 | 0 | 0 | 0 | 0 | 0 |
| | E | 0 | 0 | 0 | 0 | 0 | 0 | 0 | 0.0050 | 0 | 0 | 0 | 0 | 0 | 0 | 0 | 0 | 0 | 0 | 0 | 0 |
| | G | 0.0726 | 0 | 0 | 0.0033 | 0.0004 | 0 | 0 | 6.1655 | 0 | 0 | 0 | 0 | 0 | 0 | 0 | 0.0054 | 0 | 0 | 0 | 0.0012 |
| | H | 0.0021 | 0 | 0 | 0 | 0 | 0 | 0 | 0 | 0 | 0 | 0 | 0 | 0 | 0 | 0 | 0 | 0 | 0 | 0 | 0 |
| | I | 0 | 0 | 0 | 0 | 0 | 0 | 0 | 0 | 0 | 0 | 0 | 0 | 0 | 0 | 0 | 0 | 0 | 0 | 0 | 0 |
| | L | 0.0012 | 0 | 0 | 0 | 0 | 0 | 0 | 0.0008 | 0 | 0 | 0 | 0 | 0 | 0 | 0 | 0 | 0 | 0 | 0 | 0 |
| | K | 0 | 0 | 0 | 0 | 0 | 0 | 0 | 0.0017 | 0 | 0 | 0 | 0 | 0 | 0 | 0 | 0.0553 | 0 | 0 | 0 | 0 |
| | M | 0 | 0 | 0 | 0 | 0 | 0 | 0 | 0.0008 | 0 | 0 | 0 | 0 | 0 | 0 | 0 | 0 | 0 | 0 | 0 | 0 |
| | F | 0 | 0 | 0 | 0 | 0 | 0 | 0 | 0 | 0 | 0 | 0 | 0 | 0 | 0 | 0 | 0 | 0 | 0 | 0 | 0 |
| | P | 0.0008 | 0 | 0 | 0 | 0 | 0 | 0 | 0.0128 | 0 | 0 | 0 | 0 | 0 | 0 | 0 | 0 | 0 | 0 | 0 | 0 |
| | S | 0.0933 | 0 | 0 | 0.0004 | 0 | 0 | 0.0004 | 6.2679 | 0 | 0 | 0 | 0 | 0 | 0 | 0 | 0.0045 | 0 | 0 | 0 | 0.0025 |
| | T | 0.0004 | 0 | 0 | 0 | 0 | 0 | 0 | 0.0041 | 0 | 0 | 0 | 0 | 0 | 0 | 0 | 0 | 0 | 0 | 0 | 0 |

| | | | | | | | | | | | | | | | | | | | | | |
|---|---|---|---|---|---|---|---|---|---|---|---|---|---|---|---|---|---|---|---|---|---|
| | W | 0.0004 | 0 | 0 | 0 | 0 | 0 | 0 | 0.0008 | 0 | 0 | 0 | 0 | 0 | 0 | 0 | 0 | 0 | 0 | 0 | 0 |
| | Y | 0 | 0 | 0 | 0 | 0 | 0 | 0 | 0.0012 | 0 | 0 | 0 | 0 | 0 | 0 | 0 | 0 | 0 | 0 | 0 | 0 |
| | V | 0.0062 | 0 | 0 | 0 | 0 | 0 | 0.0004 | 0.7623 | 0 | 0 | 0 | 0 | 0 | 0 | 0 | 0.0008 | 0 | 0 | 0 | 0 |
| **NNK2 Q132X** | | 1.7513 | 0.1094 | 0.1461 | 0.0466 | 7.3546 | X | 0.0037 | 0.4222 | 0.0301 | 0.3521 | 18.8998 | 0.0029 | 0.9485 | 3.2177 | 10.9591 | 0.1255 | 0.0900 | 0.4573 | 0.0277 | 8.0575 |
| **NNK3 D138X** | | 0.0008 | 0 | 0.0017 | X | 0 | 0 | 0.0021 | 0.0140 | 0.0004 | 0 | 0 | 0 | 0 | 0 | 0 | 0 | 0 | 0 | 0 | 0.0012 |
| **NNK4 A139X** | | X | 19.5870 | 0.0012 | 0.0008 | 0.0017 | 0.0017 | 0.0041 | 8.5252 | 0.0012 | 0 | 0.0008 | 0.0025 | 0.0037 | 0.0598 | 0.2055 | 0.0054 | 0.0025 | 0.0037 | 0.3735 | 0.0062 |

### Third drug-sensitivity selection; sort 6 - population V5-low

| | | A | R | N | D | C | Q | E | G | H | I | L | K | M | F | P | S | T | W | Y | V |
|---|---|---|---|---|---|---|---|---|---|---|---|---|---|---|---|---|---|---|---|---|---|
| **NNK1 A130X R134X** | A | 0.0095 | 0 | 0 | 0 | 0 | 0 | 0.0003 | 0.1447 | 0 | 0 | 0 | 0 | 0 | 0 | 0 | 0.0008 | 0 | 0 | 0 | 0 |
| | R | X | 0 | 0 | 0.0008 | 0 | 0 | 1.4730 | 0.0767 | 0 | 0 | 0 | 0 | 0 | 0 | 0 | 0.0003 | 0.0003 | 0 | 0 | 0 |
| | N | 0 | 0 | 0 | 0 | 0 | 0 | 0 | 0 | 0 | 0 | 0 | 0 | 0 | 0 | 0 | 0 | 0 | 0 | 0 | 0 |
| | D | 0 | 0 | 0 | 0 | 0 | 0 | 0 | 0 | 0 | 0 | 0 | 0 | 0 | 0 | 0 | 0 | 0 | 0 | 0 | 0 |
| | C | 0.0005 | 0 | 0 | 0 | 0 | 0 | 0.0015 | 0.0003 | 0 | 0 | 0 | 0 | 0 | 0 | 0 | 0 | 0 | 0 | 0 | 0 |
| | Q | 0 | 0 | 0 | 0 | 0 | 0 | 0 | 0 | 0 | 0 | 0 | 0 | 0 | 0 | 0 | 0 | 0 | 0 | 0 | 0 |
| | E | 0 | 0 | 0 | 0 | 0 | 0 | 0 | 0.0061 | 0 | 0 | 0 | 0 | 0 | 0 | 0 | 0 | 0 | 0 | 0 | 0 |
| | G | 0.0025 | 0 | 0 | 0.0005 | 0 | 0 | 0.0005 | 0.3610 | 0 | 0 | 0 | 0 | 0 | 0 | 0 | 0.0018 | 0 | 0 | 0 | 0.0005 |
| | H | 0.0003 | 0 | 0 | 0 | 0 | 0 | 0.0010 | 0 | 0 | 0 | 0 | 0 | 0 | 0 | 0 | 0 | 0 | 0 | 0 | 0 |
| | I | 0 | 0 | 0 | 0 | 0 | 0 | 0 | 0 | 0 | 0 | 0 | 0 | 0 | 0 | 0 | 0 | 0 | 0 | 0 | 0 |
| | L | 0.0030 | 0 | 0 | 0 | 0 | 0 | 0.0013 | 0.0127 | 0 | 0 | 0 | 0 | 0 | 0 | 0 | 0 | 0 | 0 | 0 | 0 |
| | K | 0 | 0 | 0 | 0 | 0 | 0 | 0 | 0 | 0 | 0 | 0 | 0 | 0 | 0 | 0 | 0 | 0 | 0 | 0 | 0 |
| | M | 0.0003 | 0 | 0 | 0.0003 | 0 | 0 | 0 | 0.0386 | 0 | 0 | 0 | 0 | 0 | 0 | 0 | 0 | 0 | 0 | 0 | 0 |
| | F | 0 | 0 | 0 | 0 | 0 | 0 | 0 | 0 | 0 | 0 | 0 | 0 | 0 | 0 | 0 | 0 | 0 | 0 | 0 | 0 |
| | P | 0 | 0 | 0 | 0 | 0 | 0 | 0.0008 | 0.0008 | 0 | 0 | 0 | 0 | 0 | 0 | 0 | 0 | 0 | 0 | 0 | 0 |
| | S | 0.0020 | 0 | 0 | 0 | 0 | 0 | 0.0003 | 0.1764 | 0 | 0 | 0 | 0 | 0 | 0 | 0 | 0 | 0 | 0 | 0 | 0.0003 |
| | T | 0 | 0 | 0 | 0 | 0 | 0 | 0 | 0 | 0 | 0 | 0 | 0 | 0 | 0 | 0 | 0 | 0 | 0 | 0 | 0 |
| | W | 0 | 0 | 0 | 0 | 0 | 0 | 0 | 0 | 0 | 0 | 0 | 0 | 0 | 0 | 0 | 0 | 0 | 0 | 0 | 0 |
| | Y | 0 | 0 | 0 | 0 | 0 | 0 | 0 | 0 | 0 | 0 | 0 | 0 | 0 | 0 | 0 | 0 | 0 | 0 | 0 | 0 |
| | V | 0.4923 | 0.0010 | 0 | 0.0312 | 0.0038 | 0 | 0.0427 | 71.8158 | 0 | 0 | 0 | 0 | 0 | 0 | 0 | 0.0350 | 0.0005 | 0 | 0 | 0.0033 |
| **NNK2 Q132X** | | 0.0307 | 0.0030 | 0.0003 | 0 | 0.1170 | X | 0.0003 | 0.0206 | 0.0003 | 0.0536 | 3.3018 | 0 | 0.0498 | 0.0724 | 0.1203 | 0.0310 | 0.0005 | 0.0053 | 0.0008 | 0.1858 |
| **NNK3 D138X** | | 0 | 0 | 0.0008 | X | 0 | 0 | 0 | 0.0028 | 0 | 0 | 0 | 0 | 0 | 0 | 0 | 0 | 0 | 0 | 0 | 0 |
| **NNK4 A139X** | | X | 19.9162 | 0.0008 | 0 | 0.0005 | 0.0005 | 0 | 0.2381 | 0 | 0 | 0.0003 | 0.0074 | 0.0015 | 0.0008 | 0.0013 | 0.0074 | 0.0048 | 0.0015 | 0.0416 | 0.0003 |

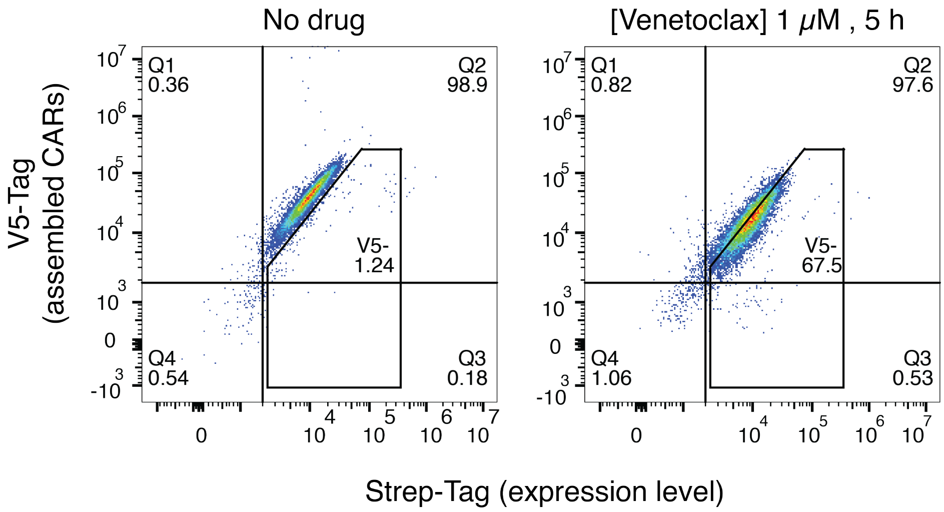


**Supplementary Figure 1: Proof-of-principle for cell-surface expression of the DROP-CAR and dissociation of the R-domain from the S-chain by venetoclax.**

Cell surface staining of B3Z cells expressing the non-optimized DROP-CAR with wild-type LD3. The V5 tag was used to stain the soluble scFv-Bcl2 and the Strep tag was used to stain the membrane bound LD3. **Left:** Tag-staining in the absence of venetoclax, **Right:** Tag-staining after 5h coincubation with 1uM venetoclax. A downwards shift of the population indicates CAR disassembly.

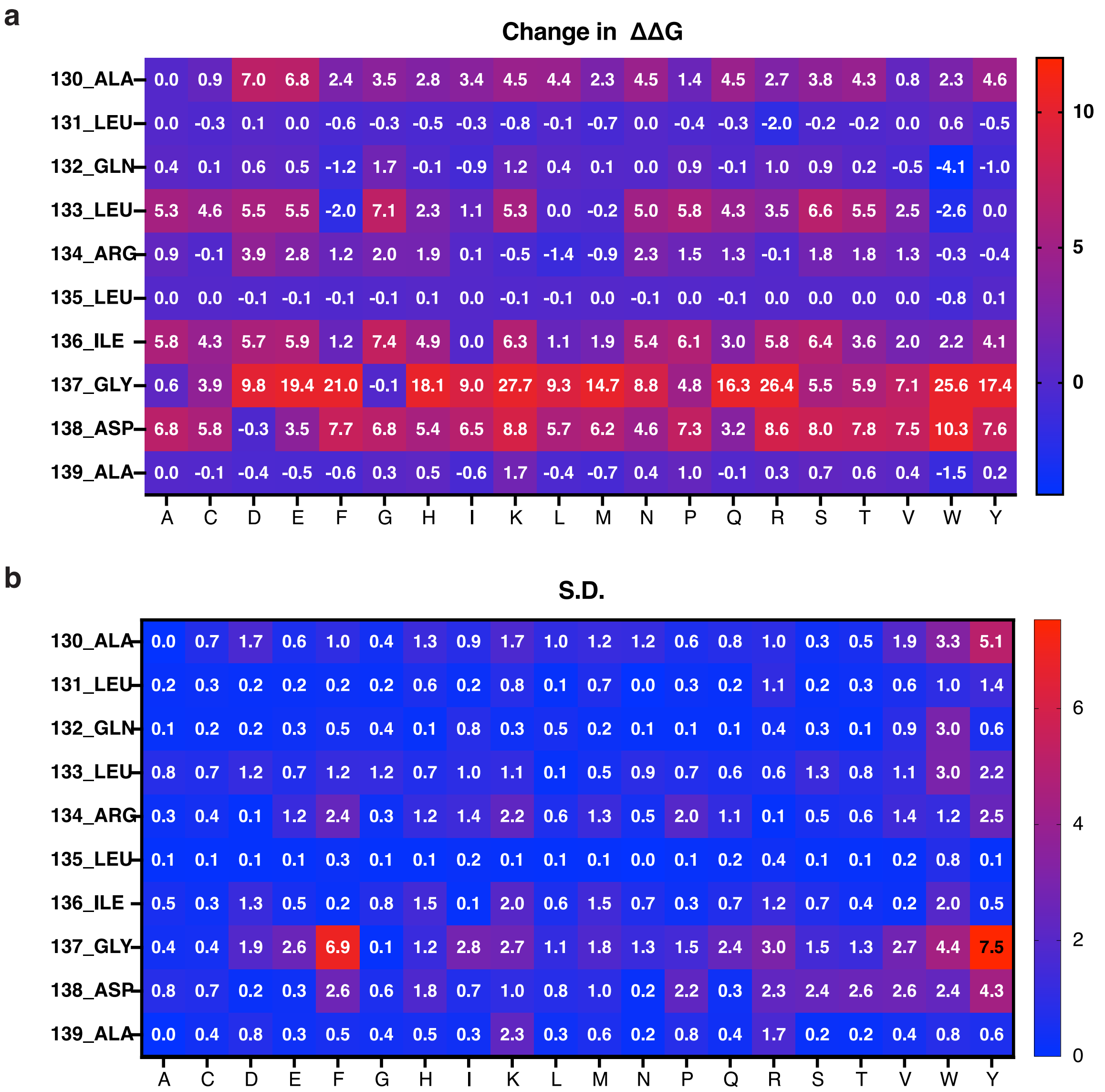


**Supplementary Figure 2: In Silico Site Saturation Mutagenesis.**

**a, b,** Heat map for the mean values and standard deviation of changes in Gibbs free energy (ΔΔG) in Rosetta energy units resulting from site saturation mutagenesis of interface residues predicted by Rosetta rigid-body docking. The residues chosen for experimental mutagenesis are shown in bold on the Y-axis, and possible amino acid replacements on the X-axis. S.D. = standard deviation.

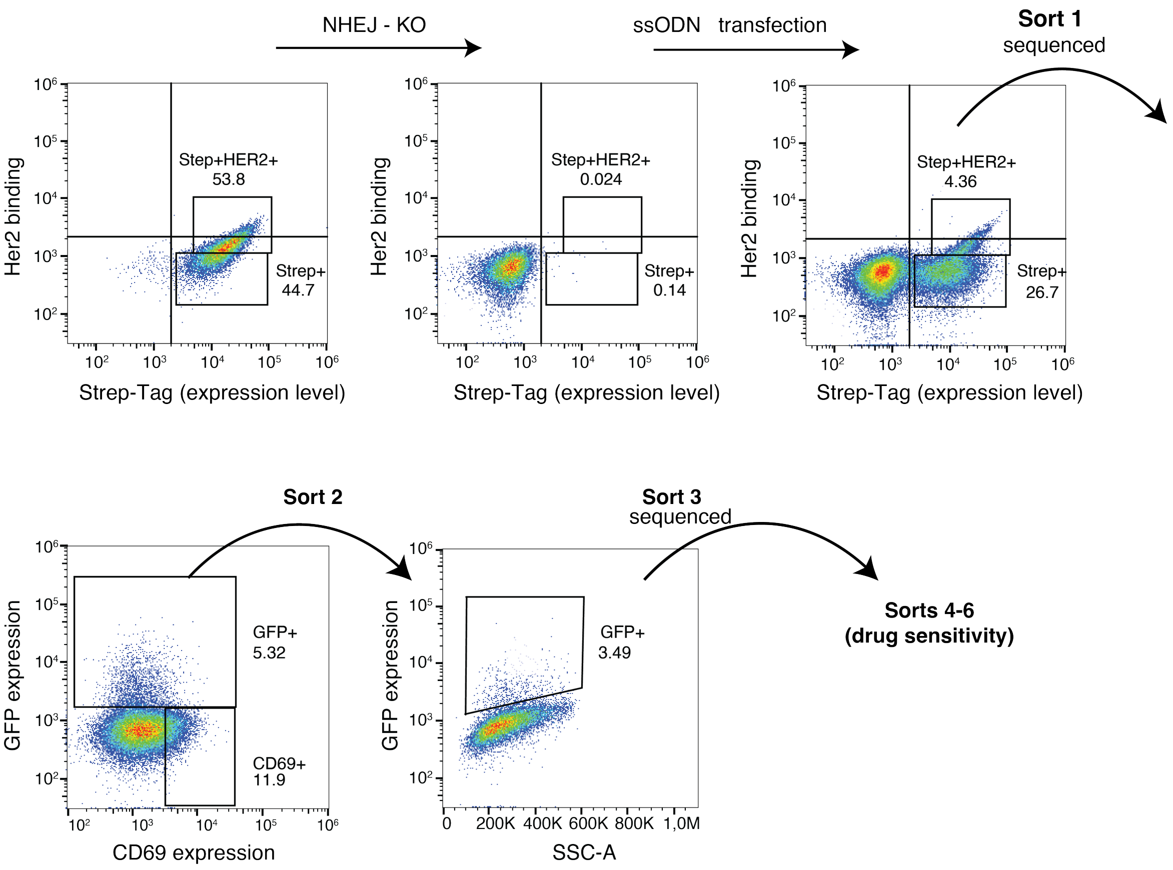


**Supplementary Figure 3: Library generation and DROP-CAR functionality sorts.** From the initial Cas9 expressing B3Z-Drop-CAR cell line, CAR expressing cells were treated with a gRNA that cuts in the LD3 region and is repaired by nonhomologous end joining (NHEJ), causing frameshift mutations. The sorted cell population without CAR expression was transfected with the ssODN library to integrate the variant library into the genomic LD3 site by homology-directed repair (HDR). The population with high expression of fully assembled DROP-CARs (costained for the V5 tag on the R-domain and the Strep tag on the transmembrane S-chain) was sorted and used for coculture experiments with Her2$^{+}$ SKOV3 cells. High hxpression of the reporter gene GFP was used to enrich cells expressing functional DROP-CARs on their surface. The functional sort was conducted twice, followed by the drug sensitivity screening with venetoclax (sorts 4-6 shown in Supplementary Figure 4). Deep sequencing results for sorts 1 and 3 are listed in Table S4.

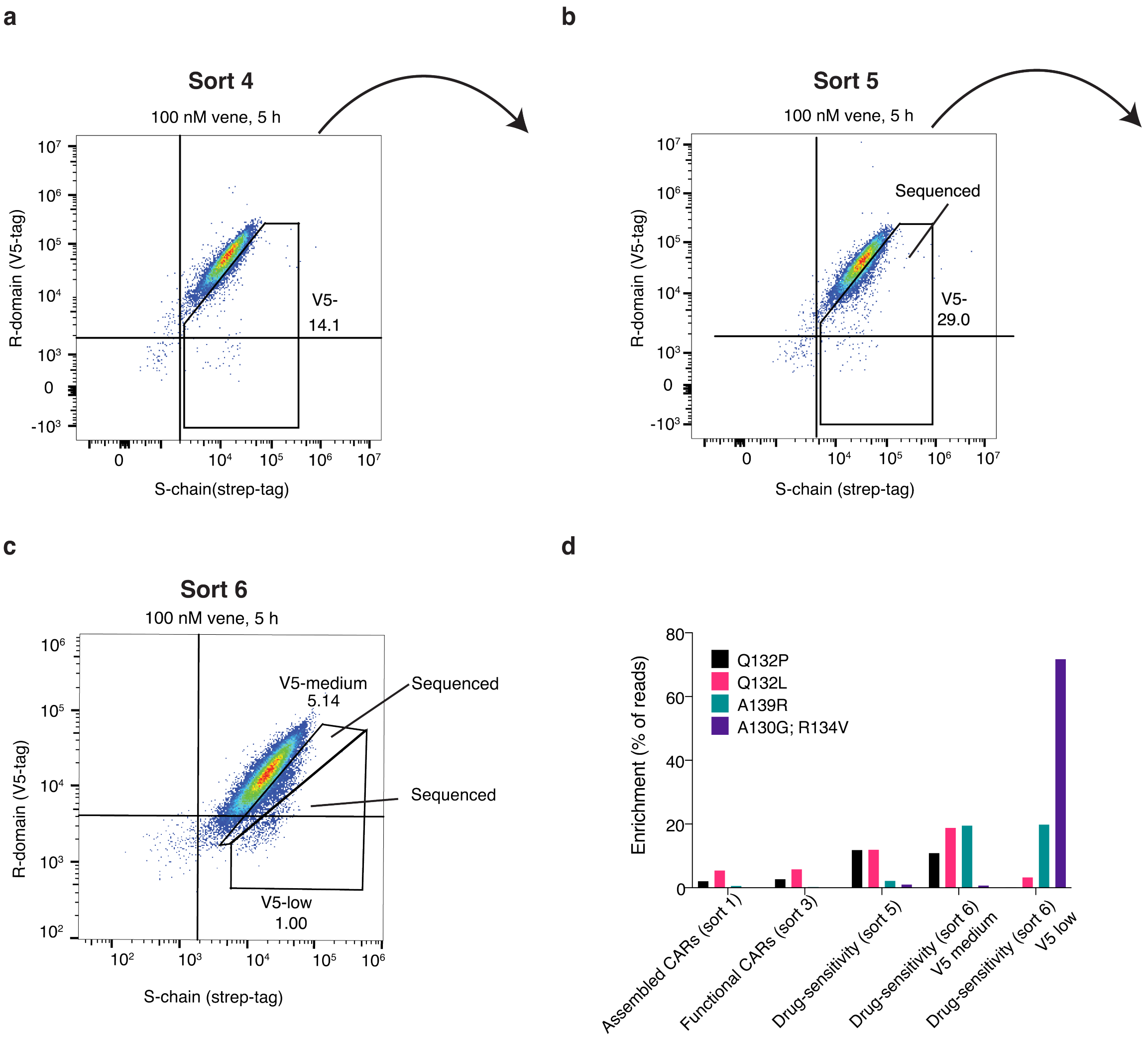


**Supplementary Figure 4: Drug sensitivity screening.**

**a-c,** Drug sensitive DROP-CARs were enriched in 3 consecutive screens (gating shown for sorted cells, sorts 4-6). Weaker V5 staining in the presence of venetoclax indicates DROP-CAR disassembly. Populations chosen for deep sequencing are indicated. (c is the same as shown in Figure 2e.) **d,** Enrichment of clones making up more than 10% of the entire population after sorts as indicated. Deep sequencing results are listed in Table S4.

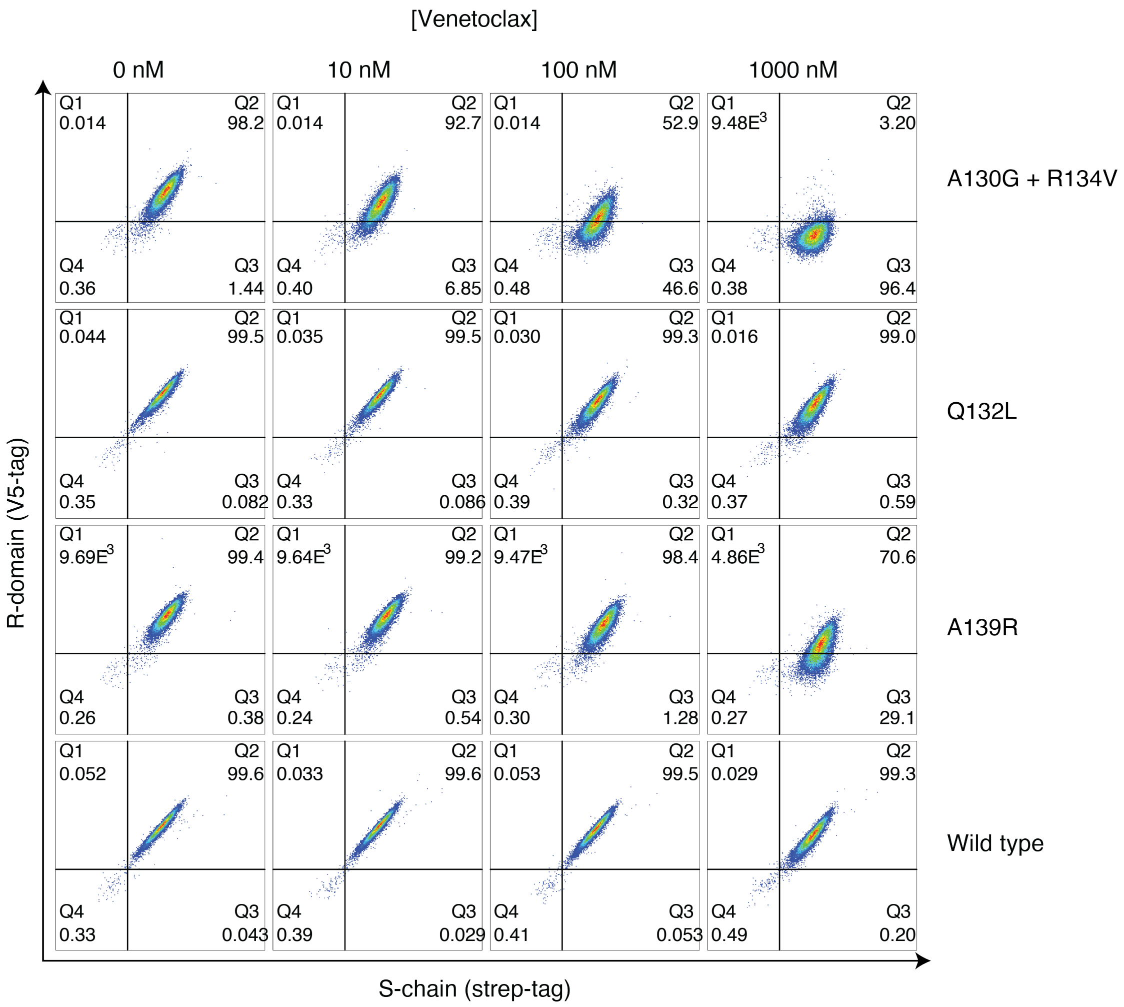


**Supplementary Figure 5: Characterization of enriched DROP-CAR (LD3) variants.**

The top hits of the enriched sequences from the V5-low and V5-medium populations of the third drug-sensitivity sort were analyzed after venetoclax treatment (5 h incubation). A decrease in the population for V5-tag staining in response to venetoclax indicates successful venetoclax-induced CAR disassembly. The first and last panels shown for the wild type LD3 are the same as in Figure S1.

```
Range 1: 23 to 166
Identities:132/144(92%), Positives:135/144(93%), Gaps:0/144(0%)

Query  1     GQRWELALGRFLEYLSWVSTLSEQVQEELLSSQVTQELRALMDETMKELKAYKSELEEQL  60
             GQRWELALGRF +YL WV TLSEQVQEELLSSQVTQELRALMDETMKELKAYKSELEEQL
Sbjct  23    GQRWELALGRFWDYLRWVQTLSEQVQEELLSSQVTQELRALMDETMKELKAYKSELEEQL  82

Query  61    TPVAEETRARLSKELQAAQARLGADMEDVRGRLVQYRGEVQAMLGQSTEELRVRLASHLI  120
             TPVAEETRARLSKELQAAQARLGADMEDVRGRLVQYRGEVQAMLGQSTEELRVRLASHL
Sbjct  83    TPVAEETRARLSKELQAAQARLGADMEDVRGRLVQYRGEVQAMLGQSTEELRVRLASHLR  142

Query  121   GLQLVLIGDAFDLQKRLAVYQAGA  144
              L+   L+ DA DLQKRLAVYQAGA
Sbjct  143   KLRKRLLRDADDLQKRLAVYQAGA  166
```

**Supplementary Figure 6: Alignment of variants with Human Apolipoprotein E4 (ApoE4).**

Alignment of the mutant LD3 A130G; R134V (Query) to human ApoE4 (Sbjct; PDB 1GS9, UniProt ID P02649) via NCBI Protein BLAST. All differences compared to the human protein are highlighted. The numbering of the mutants in the main text is based on the crystal structure PDB 6IWB, whereas the alignment starts with the first residue used in the DROP-CAR. Accordingly, the sites of the double mutant correspond to A121G; R125V in the alignment. Range 1 refers to the aligned region of ApoE4.

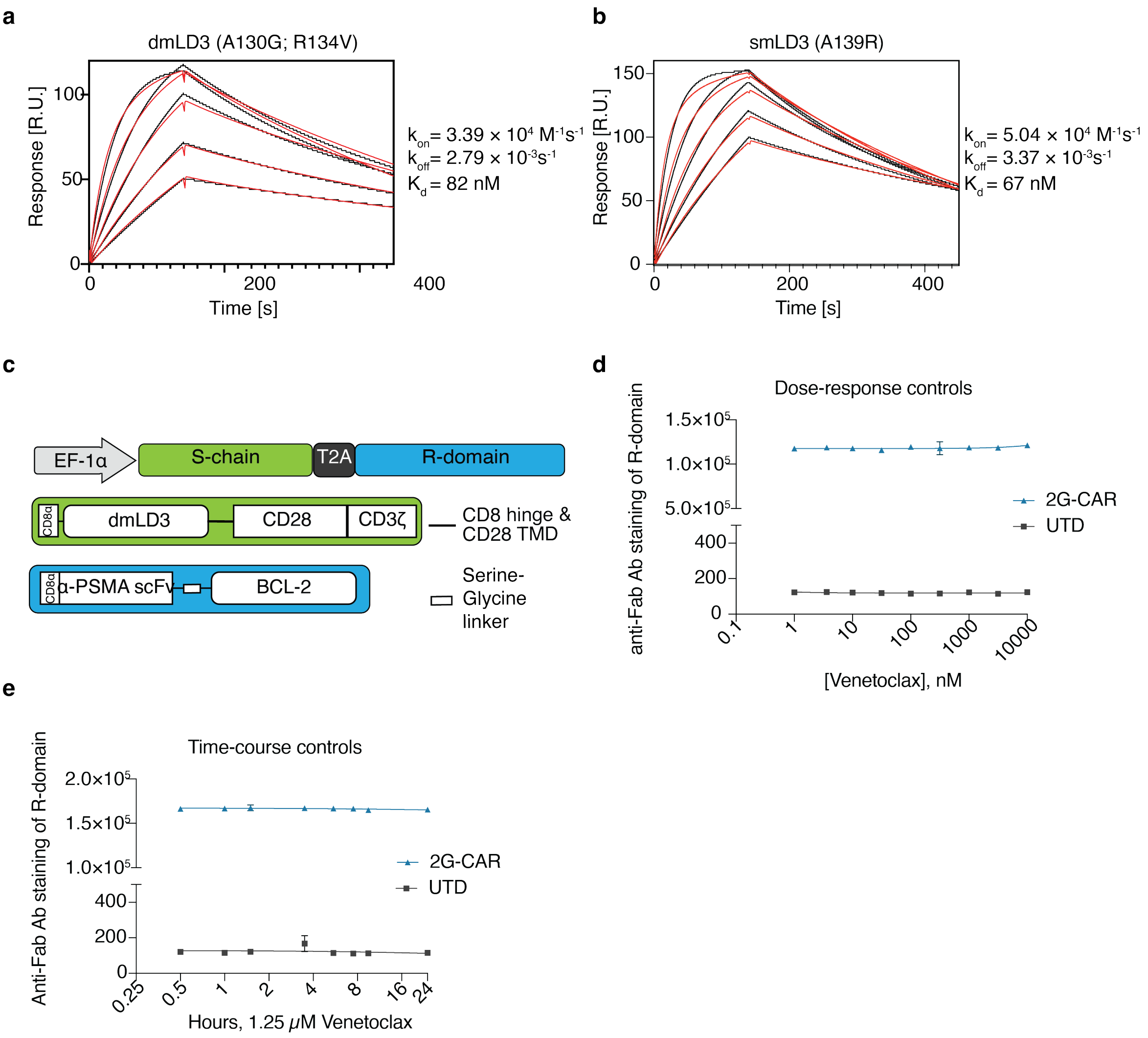


**Supplementary Figure 7: Characterization of LD3 variants enriched from the library screening and evaluation of optimized DROP-CAR expression and function in Jurkat cells.**

**a,** SPR analysis of the dmLD3 variant (A130G+R134V) and Bcl-2. **b,** SPR analysis of the single (s)LD3 variant (A139R) and Bcl-2. **c,** Schematic of the anti-PSMA DROP-CAR lentiviral expression cassette. **d,** Positive (second generation 2G-CAR) and negative (untransduced; UTD) controls for the dose-response experiment of Jurkat DROP-CAR (comprising dmLD3; A130G+R134V) disruption **e,** Positive and negative controls for the time-course experiment of DROP-CAR Jurkat (comprising dmLD3) disruption.

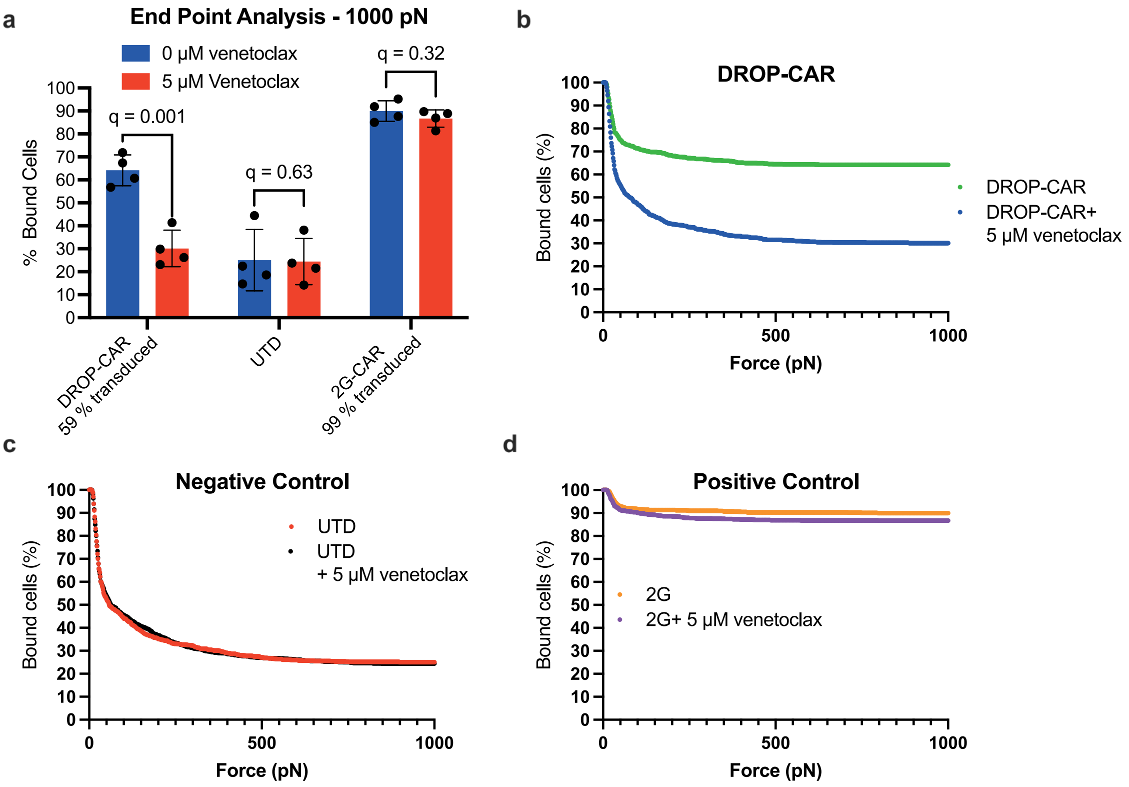


**Supplementary Figure 8: Cell avidity analysis with hinge cells counted as bound**

**a,** Percentage of cells that remain bound after 2.5 min of a linear acoustic force ramp from 0 to 1000 pN. Partly detached cells (hinge cells) are counted as bound. Values are the mean ± s.d. of n = 4 measurements on separate chips. **b,c,d,** Percent of cells bound for the DROP-Jurkat cells, untransduced cells (UTD), and positive control second generation (2G) CAR-T cells over 2.5 min of a linear acoustic force ramp from 0 to 1000 pN. Transduction efficiency for the DROP CAR was 59% and for the 2G CAR 99%. Partly detached cells (hinge cells) are counted as bound. Values are n = 4 measurements on separate chips.

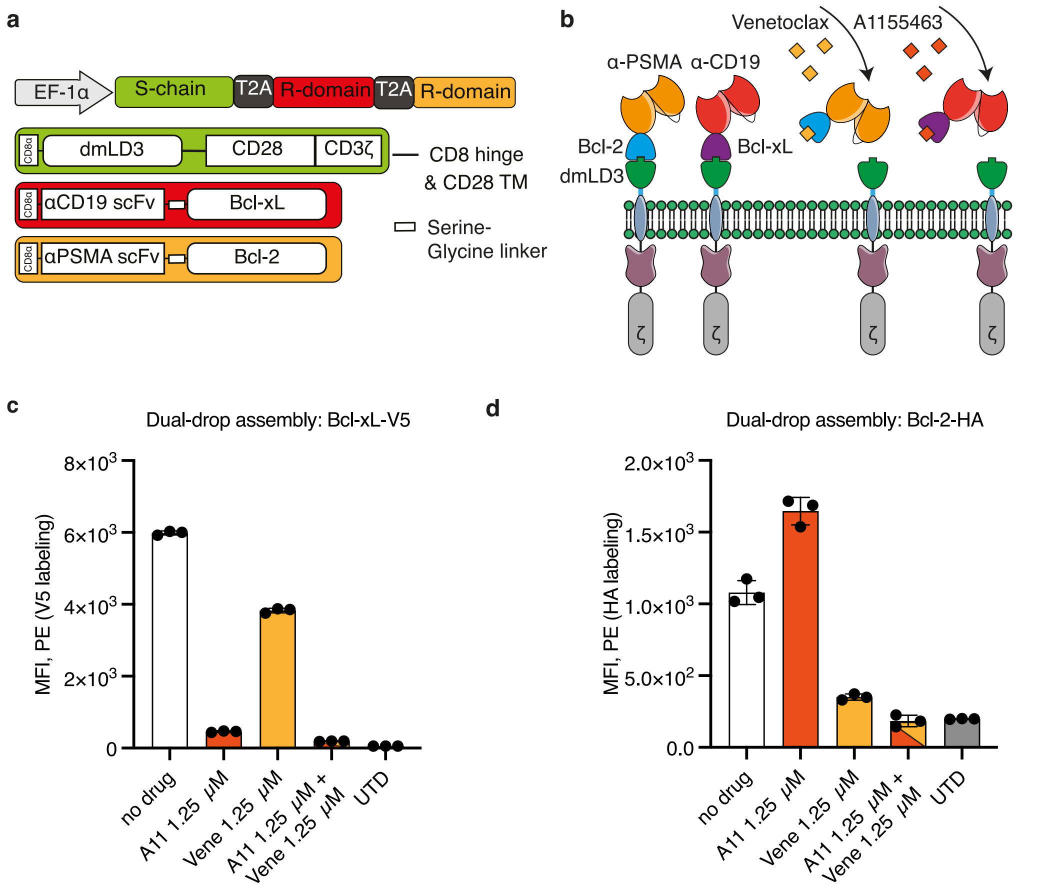


**Supplementary Figure 9: Proof of concept for dual DROP-CARs**

**a,** Schematic of a lentiviral vector encoding the dual DROP-CARs, with one R-chain comprising Bcl-xL and the other comprising Bcl-2. **b,** Schematic of cell-surface expression of two DROP-CARs, one regulated by venetoclax and the other by A1155463. **c,** Evaluation of DROP-CAR disassembly upon incubation with one or both small molecules and anti-V5 tag labeling to detect the anti-CD19 receptor domain comprising Bcl-xL. **d,** Evaluation of DROP-CAR disassembly upon incubation with one or both small molecules and anti-HA tag labeling to detect the anti-PSMA receptor domain comprising Bcl-2. Untransduced (UTD) cells were used as controls to determine background labeling. Values are the mean ± s.d. of n=3 biological replicates.

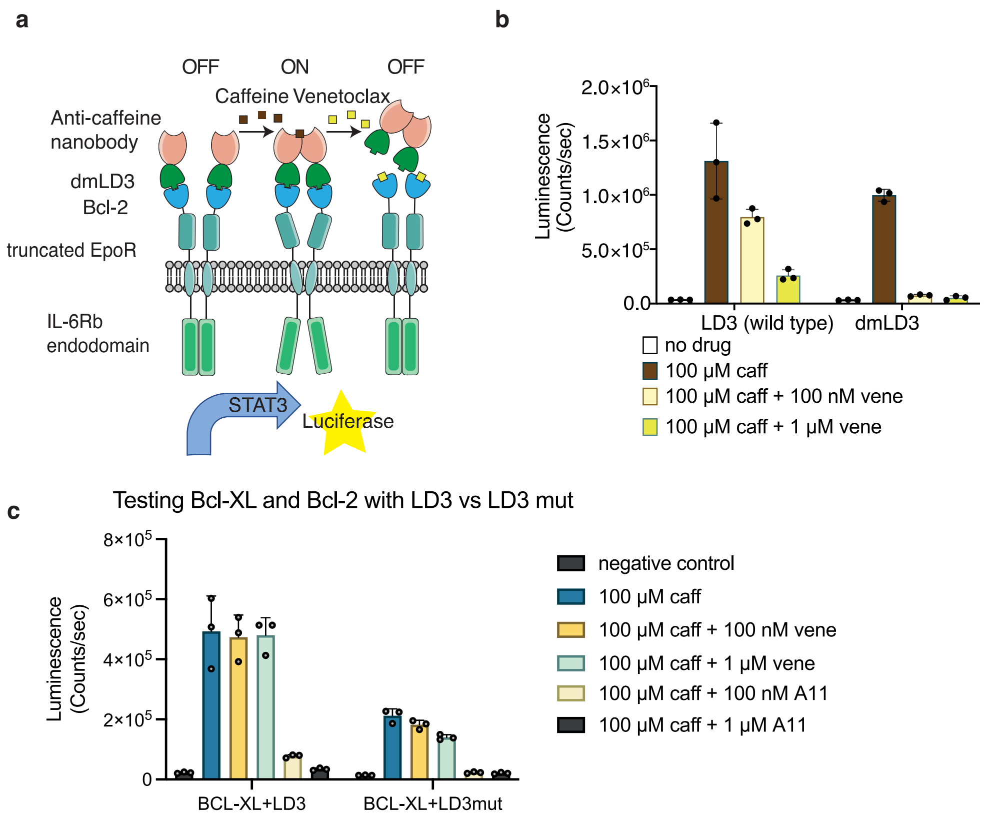


d

| A | B | A AND (NOT B) | Caffeine | Venetoclax | Caffeine AND (NOT Venetoclax) |
|---|---|---|---|---|---|
| 0 | 0 | 0 | 0 | 0 | 0 |
| 0 | 1 | 0 | 0 | 1 | 0 |
| 1 | 0 | 1 | 1 | 0 | 1 |
| 1 | 1 | 0 | 1 | 1 | 0 |

**Supplementary Figure 10: Adapting the DROP design to the generalized extracellular molecule sensor (GEMS) cytokine receptor platform.**

**a,** Schematic of engineered cytokine receptors that switch-on STAT3 signaling and STAT3-induced luciferase expression in response to caffeine and switch-off in response to Venetoclax (also shown in Figure 4a). **b,** Reporter gene expression in HEK293T cells to test Caffeine-ON, venetoclax-OFF DROP-GEMS. **c,** Reporter gene expression in HEK293T cells to test Caffeine-ON, Bcl-xL-OFF DROP-GEMS. Values are the mean ± s.d. of n=3 biological replicates.**d** Truth table for A AND (NOT B), and for caffeine AND (NOT venetoclax). This is equivalent to A NIMPLY B.

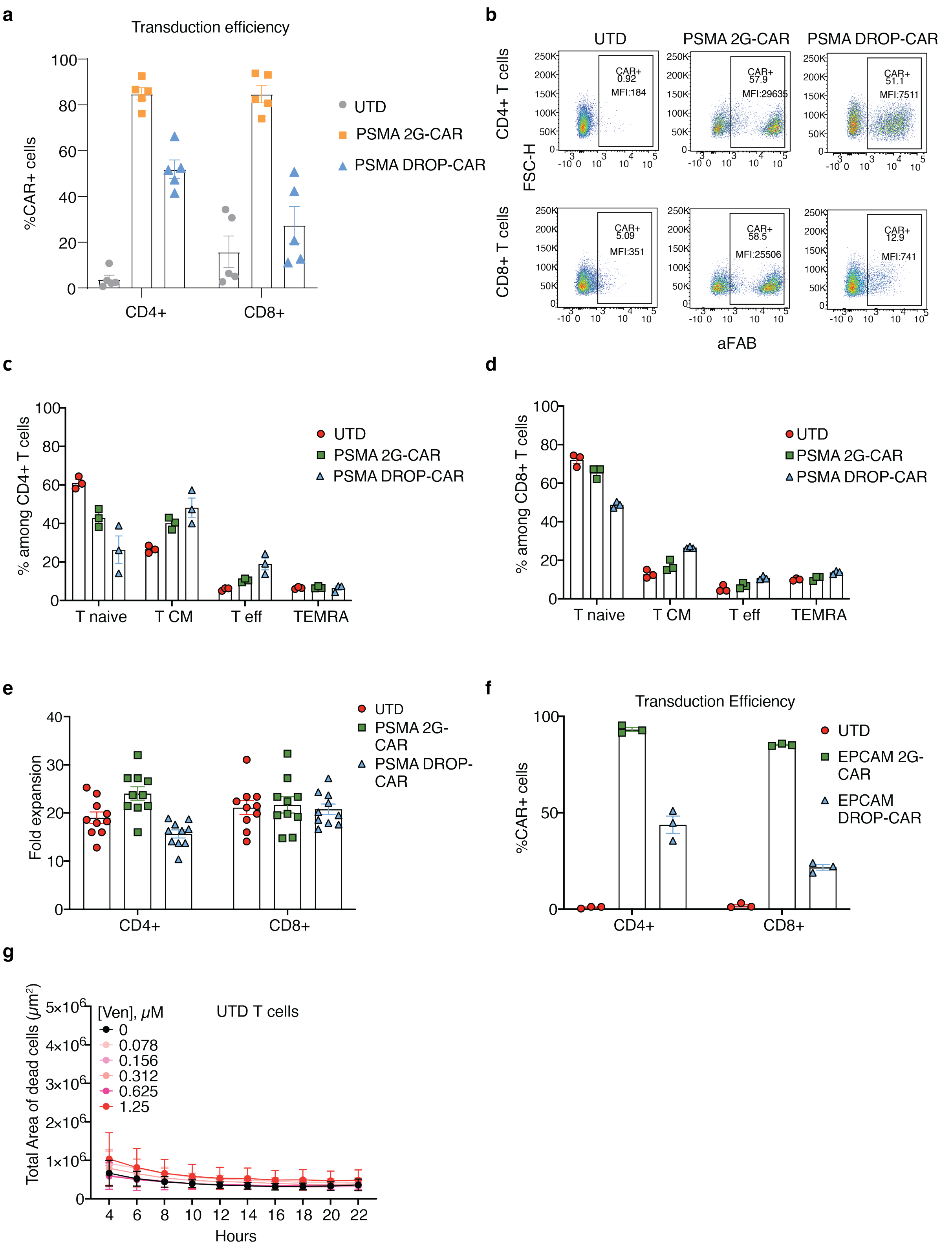


**Supplementary Figure 11: DROP-CAR expression in primary human T Cells.**

**a, b,** Anti-PSMA second generation (2G) versus DROP-CAR expression levels in CD8⁺ and CD4+ human T cells compared to untransduced (UTD) cells to determine background staining. Measured by staining the scFv of the CAR and flow cytometry.

**c,d** Phenotype ($T_{CM}$- central memory T cells, $T_{eff}$- effector T cells, $T_{EMRA}$- terminally differentiated effector memory cells re-expressing CD45RA) for human $CD4^+$ (c) and $CD8^+$ (d) T cells expressing a 2G CAR or DROP-CAR versus UTD cells. **e**, Expansion of 2G- and DROP-CAR versus UTD primary human T-cells 11 days post-transduction. **f**, Anti-EpCAM 2G versus DROP-CAR expression levels in $CD8^+$ and $CD4^+$ human T cells. **g** UTD cells as negative control for the same human donor derived cells as in Figure 5 e,f. Values are the mean ± s.e.m. of n = 3 (c,d,f,g), n = 5 (a), n = 10 (e) human donors.

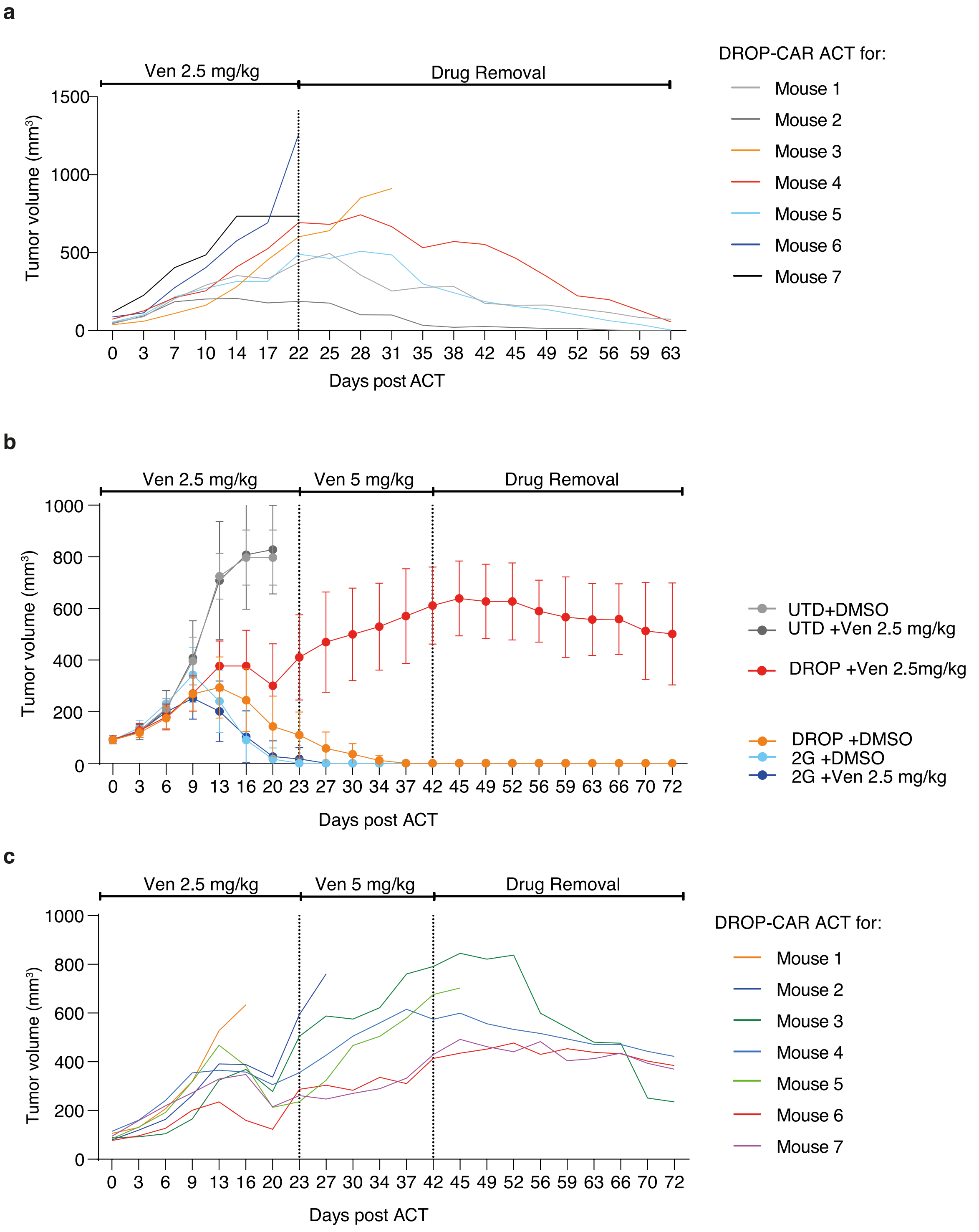


**Supplementary Figure 12: Tumor control by DROP-CAR T cells.**
**a,** Tumor growth curves of individual mice are shown for the DROP-CAR + venetoclax experimental group, before and after drug removal (Day 22). **b,** Replicate of the in vivo experiment for DROP-CAR T cells. In this experiment, at day 20 post ACT we observed incomplete repression of DROP-CAR activity in the DROP-CAR + venetoclax group compared to the vehicle control and increased the dose of venetoclax from 2.5 to 5 mg/kg. We then

ceased the venetoclax administration to monitor DROP-CAR T cell re-activation after 42 days of venetoclax exposure (this is 20 days longer compared to the 22 days of exposure in a). The graph presents the mean tumor volumes of 7 mice per group +/- s.d. **c,** Tumor growth of individual mice shown for the DROP-CAR + venetoclax experimental group for panel b.
